# Uncovering functional lncRNAs by scRNA-seq with ELATUS

**DOI:** 10.1101/2024.01.26.577344

**Authors:** Enrique Goñi, Aina Maria Mas, Amaya Abad, Marta Santisteban, Puri Fortes, Maite Huarte, Mikel Hernaez

**Affiliations:** Center for Applied Medical Research, University of Navarra, PIO XII 55 Ave, 31008, Pamplona, Spain; Institute of Health Research of Navarra (IdiSNA), Pamplona, Spain; Department of Medical Oncology, Clinica Universidad de Navarra, Pio XII 36 Ave, 31008 Pamplona, Spain; Liver and Digestive Diseases Networking Biomedical Research Centre (CIBERehd), Spanish Network for Advanced Therapies (TERAV ISCIII), Madrid, Spain; Cancer Center Clinica Universidad de Navarra (CCUN), Madrid, Spain; Data Science and Artificial Intelligence Institute (DATAI), Universidad de Navarra, Pamplona, Spain

## Abstract

Long non-coding RNAs (lncRNAs) play fundamental roles in cellular processes and pathologies, regulating gene expression at multiple levels. Despite being highly cell type-specific, their study at single-cell (sc) level has been challenging due to their less accurate annotation and low expression compared to protein-coding genes. To identify the important, albeit widely overlooked, specific lncRNAs from scRNA-seq data, here, we develop a computational framework, ELATUS, based on the pseudoaligner Kallisto that enhances the detection of functional lncRNAs previously undetected and exhibits higher concordance with the ATAC-seq profiles in single-cell multiome data. Importantly, we then independently confirmed the expression patterns of cell type-specific lncRNAs exclusively detected with ELATUS and unveiled biologically important lncRNAs, such as *AL121895.1*, a previously undocumented cis-repressor lncRNA, whose role in breast cancer progression was unnoticed by traditional methodologies. Our results emphasize the necessity for an alternative scRNA-seq workflow tailored to lncRNAs that sheds light on the multifaceted roles of lncRNAs.

## Main

Organismal functions are ultimately driven by the orchestration of the transcriptional programs of each of the individual cells that compose their tissues. The profound understanding of the cellular transcriptional configurations allows uncovering the mechanisms underlying pathological processes. While new technologies enable profiling of transcriptomes at single-cell resolution, novel computational methods are crucially needed for exploring transcriptional events at non-coding regions.

Most gene expression studies at single-cell level are exclusively focused on protein-coding genes while non-coding RNA species are very poorly investigated^1–4^. A significant fraction of non-coding RNAs are classified as long non-coding RNAs (lncRNAs), RNA Pol II transcripts lacking protein-coding potential, recently re-defined based on their length of more than 500 nucleotides^5,6^. LncRNAs are distinctively characterized by their high tissue^7^ and cell type specificity^8–10^ compared to protein-coding genes, which is linked to their regulatory functions. In line with their roles in gene regulation, alterations in lncRNA expression are associated with multiple pathologies^11,12^. All these characteristics evidence the potential benefits from their study at single-cell resolution, which is needed to achieve an improved and complete definition of cellular identity. However, limitations such as their low expression and low accuracy of their annotation have greatly hindered their use in this type of studies. According to the conservative and widely used GENCODE annotation, more than 19,000 lncRNAs are present in the human genome^13^. In contrast to the steady quantification of protein-coding genes, the annotation of lncRNAs has been in continuous evolution and growth over the last decade (**Extended Data Fig. 1**). Besides being less stable, the weak conservation levels of lncRNAs during evolution^14^ and their low expression values, complicates their detection in bulk transcriptomic data and makes the mapping of lncRNAs more challenging^15^, highlighting the need for appropriate computational methods.

Single-cell RNA-sequencing (scRNA-seq) has transformed transcriptomics by enabling the investigation of gene expression in individual cells, providing a comprehensive characterization of tissues^16–18^ and allowing the inspection of cell dynamics^19^. Particularly, scRNA-seq droplet-based methods^20,21^, predominantly the 10x Genomics technology, have revolutionized the procedure by increasing the throughput of cells and decreasing the sequencing costs^22–24^.

The computational pipeline of 10x Genomics scRNA-seq experiments begins with a preprocessing step of the sequenced samples to generate the unfiltered cell-by-gene count matrix^25^, which precedes the downstream analysis^26,27^. This critical step involves mapping the reads containing the sequenced cDNAs, as well as correcting both cell and UMI barcodes, in order to identify individual RNA molecules. Different programs based on the aligner STAR^28^, such as the widely used Cell Ranger^22^ (developed by 10x Genomics) or STARsolo^29^, perform the entire preprocessing step. In addition, the pseudoaligners Kallisto and Salmon, which are based on matching read k-mers to the transcriptome^30,31^, have also incorporated a suite of tools to preprocess the scRNA-seq sequenced reads, named Bustools^32,33^ and Alevin^34^, respectively.

Despite the differential characteristics of lncRNAs, an exhaustive comparison of scRNA-seq preprocessing pipelines focused on lncRNAs detection and quantification is missing ^27,32,34–37^. This comparison is essential to understand their underlying mechanisms and roles at single cell resolution.

Here, after benchmarking the main scRNA-seq preprocessing alternatives, including a computational validation and a comprehensive characterization of their divergences, we observed that Kallisto highlights in the detection and quantification of lncRNAs. Expanding on this exhaustive benchmarking, we have developed a specialized workflow, termed ELATUS, to streamline the identification of functionally relevant lncRNAs previously undetected in scRNA-seq experiments. Importantly, experimental validations identified *AL121895.1* as a *cis*-repressor specific of triple negative breast cancer cells. These results underscore ELATUS’s potential in uncovering expression patterns of cell type-specific and biologically relevant lncRNAs typically overlooked by standard scRNA-seq pipelines. Finally, the developed workflow, ELATUS is openly available as an R package to facilitate its adoption by the broader biomedical community.

## Results

### Preprocessing choices strongly affect lncRNA detection by scRNA-seq

LncRNAs constitute a very significant fraction of the cell transcriptome^38^. The annotation of lncRNAs is constantly evolving^6,15^, rendering their quantification more challenging compared to protein-coding genes (**Extended Data Fig. 1**). A tailored scRNA-seq workflow could shed light on their contribution to individual cell identity, which is largely understudied due to the technical limitations of single-cell technologies in terms of quantification depth and sparsity. We set to evaluate different steps of the scRNA-seq computational pipeline to identity the most suitable analysis for the detection of functional lncRNAs. Due to the more unprecise annotation of lncRNAs, we hypothesized that the quantification model choice could have a strong effect on their detection at single-cell level, as suggested by previous work of our group in bulk RNA-seq^39^.

We first conducted a comprehensive benchmarking of current state-of-the-art scRNA-seq preprocessing pipelines, including both the alignment-based methods Cell Ranger and STARsolo, and the pseudoalignment-based methods Kallisto-Bustools and Salmon-Alevin, to evaluate how they affect the detection and quantification of lncRNAs (**Fig. 1a**). To this end, we first used widely characterized 10x Genomics datasets: 1k brain cells from an E18 mouse^40^, and 10k healthy human peripheral blood mononuclear cells (PBMCs)^41^. These public datasets have been already applied for comparing the distinct pipelines^27,32,35,36,42^. In agreement with previous research^36,42^, Kallisto produced a slightly higher mapping rate (**Fig. 1b and Extended Data Fig. 2a**). Besides, the pseudoalignments-based methods presented the shortest running times (**Extended Data Fig. 2b and Extended Data Fig. 3a**) and were less memory-expensive (**Extended Data Fig. 2c and Extended Data Fig. 3b**).

**Figure 1.**
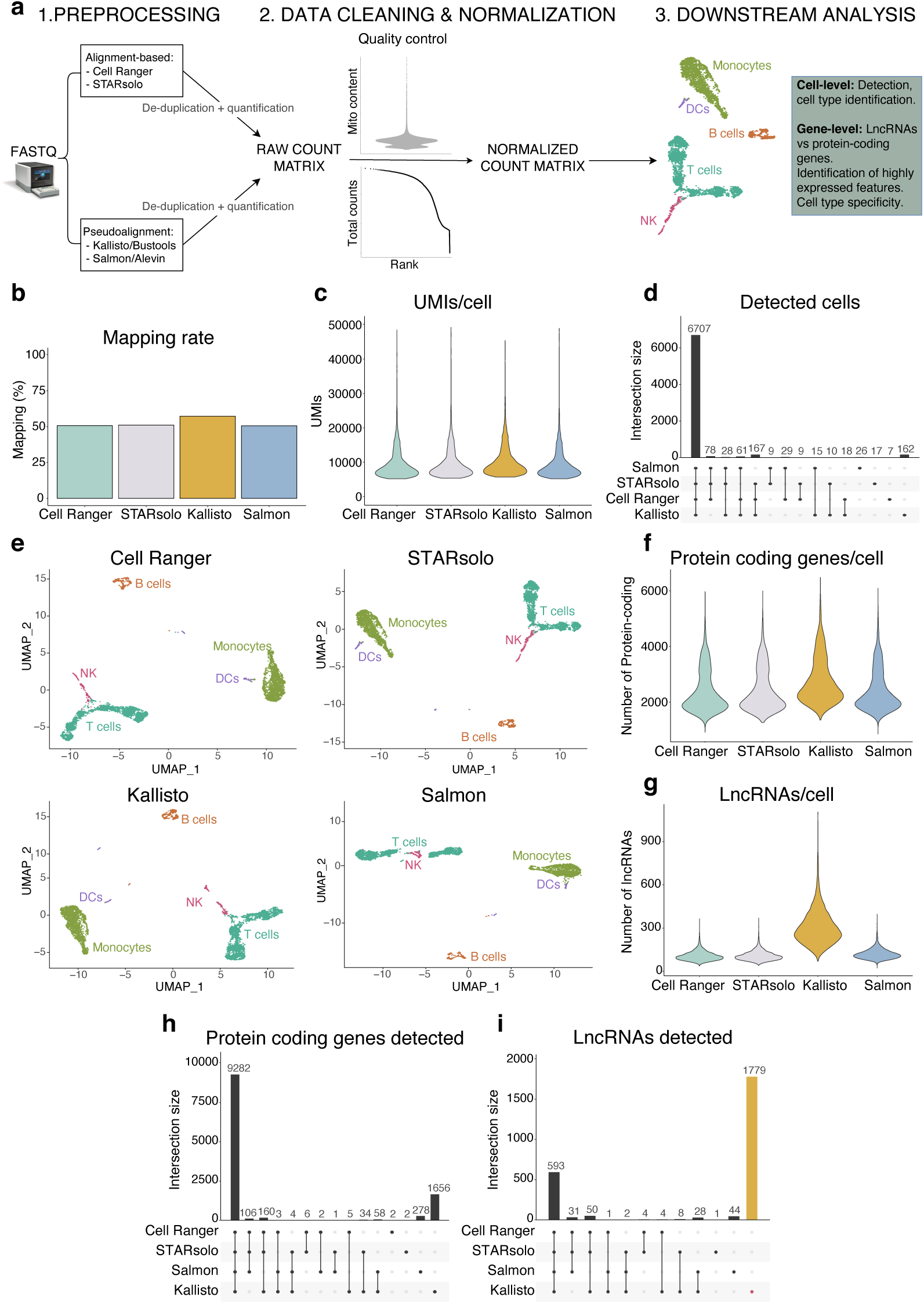
Preprocessing choices strongly affect lncRNA detection in a scRNA-seq dataset consisting of 10k human PBMCs from a healthy donor. a, Detailed step-by-step explanation of the benchmark. First, public fastq files were directly processed with the aligner-based Cell Ranger and STARsolo, as well as with the pseudoaligners Kallisto and Salmon to generate the raw count data. Next a quality control step was performed to filter empty droplets, cells with high mitochondrial content and potential multiplets, followed by a normalization step. Then, the downstream analysis began with a dimensionality reduction followed by a clustering and cell annotation steps. The detection and quantification of high-quality cells, main cell types, protein-coding genes, lncRNAs and cell type-specific features was compared depending on the preprocessing choice. **b,** Mapping rate by each pipeline. **c,** Number of UMI counts per cell by each pipeline. **d,** UpSet plot showing the overlap of retained high-quality cells by each pipeline. **e,** UMAP plots displaying the populations of the main cell types identified across pipelines. **f,** Number of detected protein-coding genes per cell across pipelines. **g,** Number of detected lncRNAs per cell across pipelines. **h,** UpSet plot displaying the overlap of highly-expressed protein-coding genes (more than 250 counts and present in more than 25 cells) per pipeline. **i,** UpSet plot displaying the overlap of highly-expressed lncRNAs (more than 250 counts and present in more than 25 cells) per pipeline.

We next conducted a quality-control step on each raw cell-by-gene UMI count matrix generated by all pipelines and filtered them for empty droplets. In human PBMCs, the mitochondrial content was very similar in all tested pipelines (**Extended Data Fig. 3c**), while it was higher with Salmon in the mouse brain dataset (**Extended Data Fig. 2d**). We then removed cells with high mitochondrial composition and potential multiplets to preserve high-quality cells^26^. We observed that these cells had comparable expression levels among pipelines (**Fig. 1c and Extended Data Fig. 2e**) and that the majority of them were commonly retained by all of them (**Fig. 1d and Extended Data Fig. 2f**). Furthermore, the main cell types were distinguished by all tested preprocessing options when using canonical markers (**Fig. 1e, Extended Data Fig. 2g-h, Extended Data Fig. 3d, Supplementary Data Table 1)**.

Interestingly, regarding gene detection, we observed important differences across pipelines. While the distribution of detected protein-coding genes per cell was more similar, the number of identified lncRNAs per cell by Kallisto was strikingly higher (**Fig. 1f-g, Extended Data Fig. 2i-j**). To exclude the possibility that these differences were caused by poorly expressed genes with practically no counts, we next retained only those that fulfilled minimal expression thresholds (see Methods). While most highly-expressed protein-coding genes were commonly detected by different pipelines (**Fig. 1h, Extended Data Fig. 2k**), a very significant number of highly-expressed lncRNAs were only recognized by Kallisto, both in human and mouse datasets, whereas the remaining pipelines did not quantify them (**Fig. 1i, Extended Data Fig. 2i**). The impact of the preprocessing choice on the detection of lncRNAs was consistent across an increasing set of thresholds on expression (**Extended Data Fig. 3e-f**).

To further investigate whether the observed differences were maintained across different models and tissues, we expanded the benchmarking and analyzed a large and diverse set of public 10x Genomics scRNA-seq datasets. Specifically, we used data from human healthy intestine^43^, healthy lung and pulmonary fibrotic samples^44,45^, as well as PBMCs from mouse^46^ and human (5k cells)^47^ (**Extended Data Fig. 4**). Due to the similarities between the results yielded by the gold-standard pipeline Cell Ranger and the other preprocessing alternatives, with the exception of Kallisto, in what follows, we restricted the assessment to these two preprocessing pipelines.

The results on the extended benchmark verified that, after filtering low-quality cells and multiplets, the expression per cell was practically identical across datasets between Cell Ranger and Kallisto (**Extended Data Fig. 4a**) and most high-quality cells were commonly retained (**Extended Data Fig. 4b**). With respect to gene detection, while the distribution of protein-coding genes detected per cell was very similar in both pipelines, we confirmed that Kallisto found a remarkably higher number of lncRNAs in each cell (**Extended Data Fig. 4c-d**). Further, using a gradient of thresholds on expression to filter poorly expressed genes, we corroborated that most protein-coding genes were commonly identified across distinct datasets, whereas there was an important fraction of lncRNAs exclusively captured with Kallisto (**Extended Data Fig. 4e-f**).

Altogether, these results indicate that, while the detection of mRNAs is not affected, the identification of lncRNAs in scRNA-seq data is severely influenced by the preprocessing choice. In particular, the Kallisto preprocessing pipeline stands out in the detection and quantification of lncRNAs in an expanded and diverse set of scRNA-seq datasets.

### scATAC-seq multiome indicates an optimized preprocessing alternative for lncRNA quantification

To assess the biological plausibility of the lncRNAs exclusively quantified by Kallisto, we employed single-cell multiome data. This dataset allows for simultaneous measurement of gene expression via RNA-seq and mapping of open-chromatin using with ATAC-seq within the same cell. We reasoned that scATAC-seq would mirror the expression of lncRNAs identified by scRNA-seq without encountering the same technical biases. Specifically, we selected a public 10x Genomics multiome dataset containing 3k PBMCs from a healthy donor^48^ and we tested whether there was more consistency between scATAC-seq profiles and scRNA-seq measurements when the latter was preprocessed with Cell Ranger or Kallisto.

We started by applying general scATAC-seq quality control thresholds (ATAC counts, TSS enrichment and nucleosome signal) to filter low-quality nuclei (**Extended Data Fig. 5a-c**). Next, we removed nuclei with high mitochondrial content (**Extended Data Fig. 5d**) and empty droplets (see Methods). The expression per nuclei was very similar, although slightly higher with Kallisto (**Fig. 2a**). Using established markers, we were able to distinguish the main cell types with both scRNA-seq pipelines (**Fig. 2b, Extended Data Fig. 5e, Supplementary Data Table 1**). To test the concordance between the scATAC-seq signal and the scRNA-seq gene expression, we constructed a gene activity matrix by counting the scATAC-seq fragments that fall in the gene body and promoter regions. Then, for every high-quality nucleus we checked which RNA-seq quantification pipeline, Cell Ranger or Kallisto, yielded more genes having coincident ATAC-seq signal and RNA-seq expression (**Fig. 2c**, see Methods). To contemplate diverse scenarios, a gradient of thresholds on both ATAC-seq and RNA-seq was used to classify a gene as simultaneously activated (see Methods). Interestingly, for the majority of thresholds, we observed that scATAC-seq coupled to scRNA-seq analyzed by Kallisto gave a statistically significant increase in the number of genes, in each nucleus, that are simultaneously activated, compared to scATAC-seq coupled to scRNA-seq analyzed by Cell Ranger (**Fig. 2d**, p-value<0.0001).

**Figure 2.**
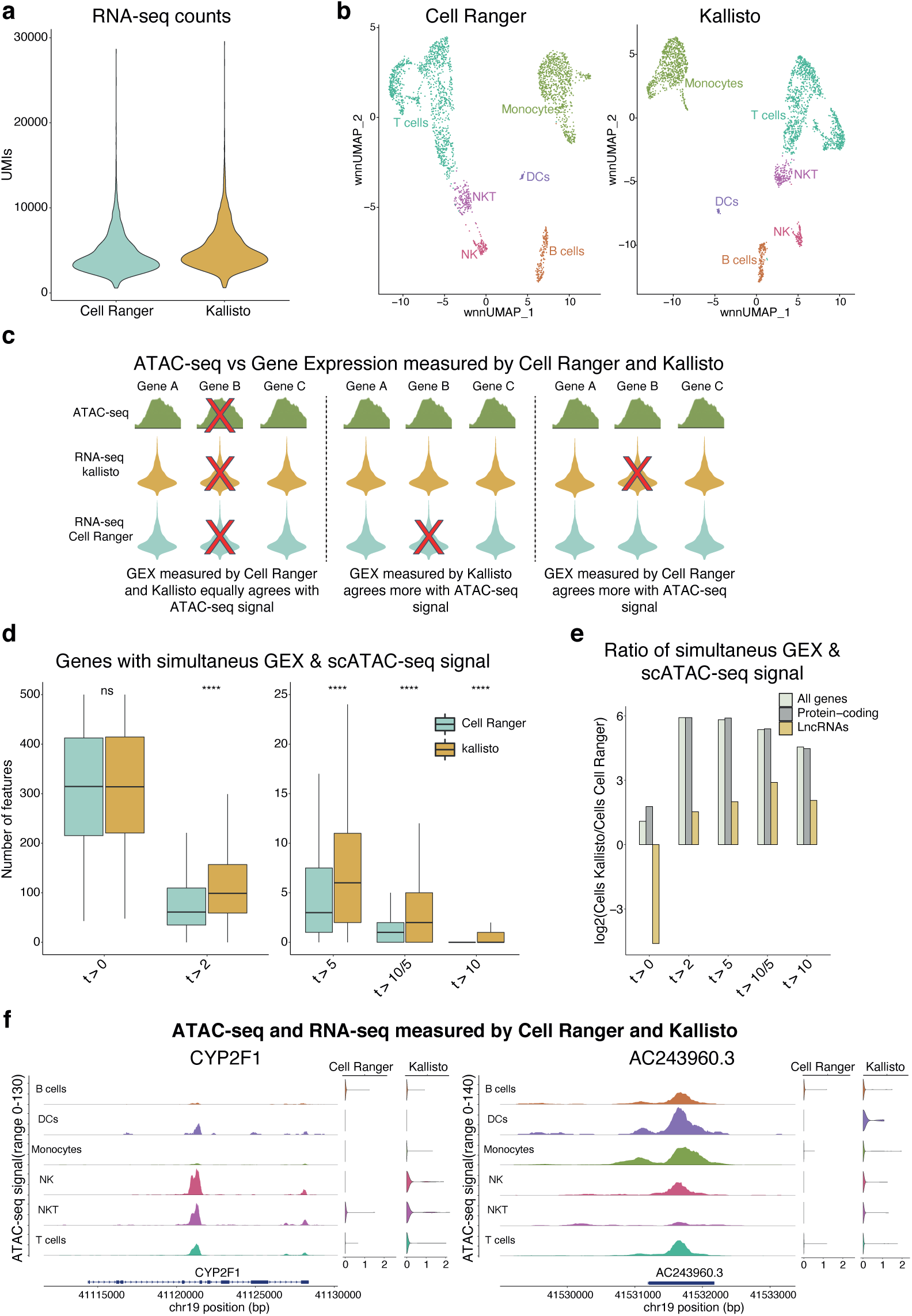
scATAC-seq multiome indicates an optimized preprocessing alternative for lncRNA quantification. **a,** Number of UMI counts per cell obtained when preprocessing the scRNA-seq with Cell Ranger and Kallisto**. b,** Weighted nearest neighbors UMAP plot displaying the populations of cell types identified by Cell Ranger and Kallisto. **c,** Methodology used for comparing the similarity between the scRNA-seq, processed with Cell Ranger or Kallisto, and the Gene Activity matrix obtained from the scATAC-seq data. Specifically, for each nucleus, we count the number of simultaneously expressed genes in the scRNA-seq (with higher expression than a threshold), both with Cell Ranger and Kallisto, and in the Gene Activity matrix (with higher signal than a threshold). **d,** Boxplot displaying, for each nucleus, the number of simultaneously activated genes when the scRNA-seq is processed with Cell Ranger and Kallisto. Two-tailed student t-test for assessing differential expression. **e,** Ratio of the number of nuclei for which there is more genes simultaneously activated with Kallisto divided by the number of nuclei for which there is more genes simultaneously activated with Cell Ranger. For each nucleus we have considered the expression of all genes (white), only protein-coding genes (grey) and only lncRNAs (yellow). In (**d**) and (**e**), the x-axis represents the different thresholds used for quantifying only a gene as simultaneously activated if it had: (t>0) at least 1 UMI in RNA-seq and 1 read in ATAC-seq and, (t>2) at least 3 UMIs in RNA-seq and 3 reads in ATAC-seq, (t>5) at least 6 UMIs in RNA-seq and 6 reads in ATAC-seq, (t>10/5) at least 11 UMIs in RNA-seq and 6 reads in ATAC-seq and (t>10) at least 11 UMIs in RNA-seq and 11 reads in ATAC-seq. **f,** ATAC-seq signal and RNA-seq expression, with both Cell Ranger and Kallisto, of protein-coding gene *CYP2F1* (left) and lncRNA *AC243960.3* (right).

Moreover, the number of nuclei that had more genes coincidently activated according to RNA-Seq and ATAC-Seq was, in general, remarkably higher for Kallisto than Cell Ranger across all thresholds (**Fig. 2e, Extended Data Fig. 5f**). The improved association between scATAC-seq and scRNA-seq was illustrated in the protein-coding gene *CYP2F1* and the lncRNA *AC242960.3*, where their ATAC-seq profiles across distinct cell types only corresponded to the RNA-seq expression when processed with Kallisto (**Fig. 2f**).

Together, these results demonstrate that the lncRNAs detected by Kallisto correspond better to the scATAC-seq measurements on the same cells, associated with open and transcriptionally active chromatin, confirming that this is an optimized alternative to improve the quantification of lncRNAs.

### Exclusive and commonly identified lncRNAs share similar characteristics

In order to build up a tailored workflow for lncRNA quantification in scRNA-seq, we first needed to delve into the extra-detected lncRNAs to understand the reasons driving their detection and thus determine whether they could be potential bona fide lncRNAs. To that end, we investigated distinct properties of genes both exclusively and commonly identified by Kallisto (for simplicity termed “exclusive” and “common” genes, respectively), focusing on their expression profiles and sequence composition. In particular, we inspected their absolute expression, length and number of exons, as well as their repeat content, k-mer distribution, and cell type specificity levels. This was conducted in each scRNA-seq dataset previously included in this work, in which we applied a gradient of thresholds on expression to filter poorly-expressed genes.

We observed that both exclusive lncRNAs and protein-coding genes were significantly less expressed than the common ones (**Fig. 3a, Extended Data Fig. 6a**). Regarding their length, we noted interesting discrepancies between lncRNAs and protein-coding genes. While exclusive protein-coding genes were significantly longer for every dataset, length differences between exclusive and common lncRNAs were not significantly different for most datasets and under varied thresholds on expression (**Fig. 3b, Extended Data Fig. 6b**).

**Figure 3.**
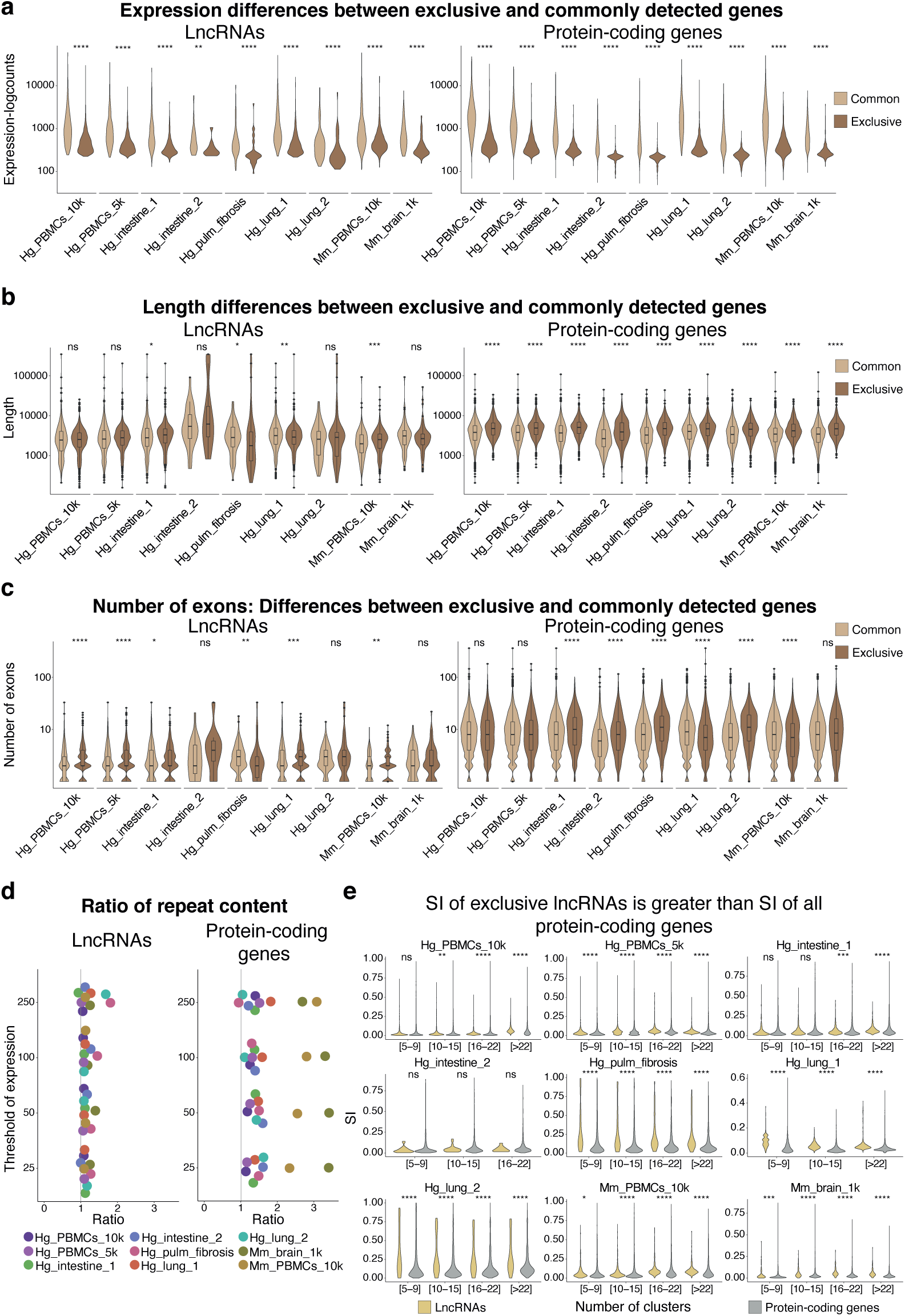
Exclusive and commonly identified lncRNAs share similar characteristics. **a,** Normalized expression differences, **b,** Length differences and **c,** Differences in the number of exons of (left) lncRNAs exclusively found by Kallisto compared to lncRNAs commonly identified by all pipelines and (right) protein-coding genes exclusively found by Kallisto compared to protein-coding genes commonly identified by all preprocessing options. In (**a**), (**b**) and (**c**) only genes with more than 250 counts and present in more than 25 cells were considered. Significance in (**a**), (**b**) and (**c**) was assessed with a two-tailed Wilcoxon test. **d,** Ratio of the sequence covered by repeats of exclusive (left) lncRNAs (right) protein-coding genes divided by the percentage of the sequence covered by repeats of common (left) lncRNAs (right) protein-coding genes. A jitter on the y-axis was included for ease of visualization. Different thresholds on expression for removing lowly-expressed genes were considered, from down to up; more than 25 counts and present in more than 3 cells, more than 50 counts and present in more than 5 cells, more than 100 counts and present in more than 10 cells and more than 250 counts and present in more than 25 cells. **e,** Specificity index (SI) differences to test if the SI of lncRNAs exclusively identified by Kallisto is significantly higher (one-tailed Wilcoxon test) than the SI of protein-coding genes. The SI distributions were calculated across distinct ranges of number and sizes of clusters, from fewer and bigger to more and smaller clusters. Starting with 5-9 clusters, then 10-15 clusters, then 16-22 clusters and finally generating a specially detailed subdivision with at least 23 clusters. Only genes with more than 250 counts and present in more than 25 cells were considered.

Given that the number of exons has been associated with the functionality of lncRNAs^10^, we wondered whether their distribution differs between exclusively detected and common genes, as an abnormal exon distribution could indicate annotation errors and therefore explain quantification differences. In general, for all different thresholds applied, the number of exons of exclusive lncRNAs was significantly higher compared to common lncRNAs. This contrasts with protein-coding genes, where there was no evident trend, since in many datasets the differences were not significant while in others significance appears in any direction (**Fig. 3c, Extended Data Fig. 7a**). The fact that exclusive lncRNAs had more exons, and hence more splicing-junctions that make them more prone to annotation inaccuracies, supported our initial hypothesis that pseudoaligners might be more suited to lncRNA quantification due to the unprecise annotation of lncRNAs^15^ compared to that of protein-coding genes.

Furthermore, it has been noted that pseudoalignments methods could cause poor quality alignments to low-complexity sequences that result in unexpected high expression of particular genes^42^. To test this possibility, we next compared the percentage of each gene that was covered by repeat elements in both exclusively detected and common features and represented this as a ratio. In agreement with this explanation, we found that repeat content of exclusive protein-coding genes was clearly larger than that of the common ones (**Fig. 3d, right**). However, the repeat content of exclusive and common lncRNAs was very similar (**Fig. 3d, left**) suggesting that the increased detection of lncRNAs by Kallisto was not caused by poor quality alignments to low-complexity sequences. In fact, the ratio between the repeat content of exclusive to common protein-coding genes was significantly higher than the ratio between the repeat content of exclusive to common lncRNAs (p-value = 4e-5, one-tailed paired t-test).

Aside, it has been documented that lncRNAs with similar k-mer profiles share related functions^49^. We applied this reasoning and analyzed the functional communities according to k-mer profiles. The analysis did not identify communities preferentially composed of exclusively captured lncRNAs. Indeed, functionally-related communities were formed by both common and exclusively identified lncRNAs (**Extended Data Fig. 7b**), suggesting that exclusive lncRNAs are not enriched in a particular function related to k-mer content but rather are comparable to known functional lncRNAs in this regard^49^.

Finally, given that lncRNAs are defined to be more cell type-specific than protein-coding genes^7–10^, we wondered whether this was maintained for the exclusive lncRNAs. Therefore, we calculated the specificity of the exclusive lncRNAs and compared it with the specificity of protein-coding genes. For this purpose, we defined a specificity index (SI, see Methods) that measured how localized or ubiquitous the expression of a gene is. To assess the influence of both the size and the number of clusters on the SI, we clustered the scRNA-seq datasets in a gradient from large to small subpopulations. In accordance with their defined properties, we corroborated that the SI exclusive lncRNAs was significantly higher than that of protein-coding genes, both in large and small subclusters in the majority of datasets (**Fig. 3e, Extended Data Fig. 7c**).

In summary, our findings indicate that exclusive lncRNAs are less expressed than common lncRNAs, but also more cell type-specific when compared to protein-coding genes. In terms of sequence composition, both groups of lncRNAs are comparable, although exclusive lncRNAs have more exons. The observations obtained from the performed characterization reinforce Kallisto’s advantages for lncRNA analysis in scRNA-seq, enabling hypothesis testing over vastly more lncRNAs than with other pipelines. However, limited by the capabilities to experimentally study lncRNAs, there is a need for specialized workflows to select lncRNAs with potential biological significance.

### Biologically relevant lncRNAs are uncovered by ELATUS

To provide a curated list of lncRNAs likely of having biological importance, we implemented a computational workflow for <u>E</u>lucidating biologically relevant lncRNAs <u>a</u>nnotated transcripts <u>u</u>sing <u>s</u>cRNA-seq, termed ELATUS. We reasoned that a biologically significant exclusive lncRNA should be expressed over a threshold and should have a highly cell type-specific expression pattern. Indeed, we observed that the majority of exclusive lncRNAs were tissue-specific, reaffirming the well-known specificity of lncRNAs^7^ (**Fig. 4a, Extended Data Fig. 8a**). However, the biological relevance of these lncRNAs remains to be determined. Interestingly, exclusive lncRNAs were enriched among lncRNAs identified by CRISPRi screenings in multiple human cell lines^10^ (p-value < 0.05, hypergeometric test) (**Supplementary Data Table 2, Extended Data Fig 9**), indicating their role in supporting cellular functions. Thus, ELATUS, besides retaining robustly expressed lncRNAs detected by both Cell Ranger and Kallisto, was designed to retain exclusive lncRNAs that were highly specific according to restrictive selection thresholds (**Fig. 4b**, see Methods). Notably, ELATUS uncovered 87 cell type-specific and highly-expressed lncRNAs, which added to the 173 lncRNAs that were also undetected by Cell Ranger and were hits in the CRISPRi screenings^10^ and to the 2,080 highly-expressed commonly detected lncRNAs (**Supplementary Data Table 3**), defined a complete collection of 2,340 lncRNAs that exhibit characteristics of functional lncRNAs in the diverse set of scRNA-seq datasets analyzed.

**Figure 4.**
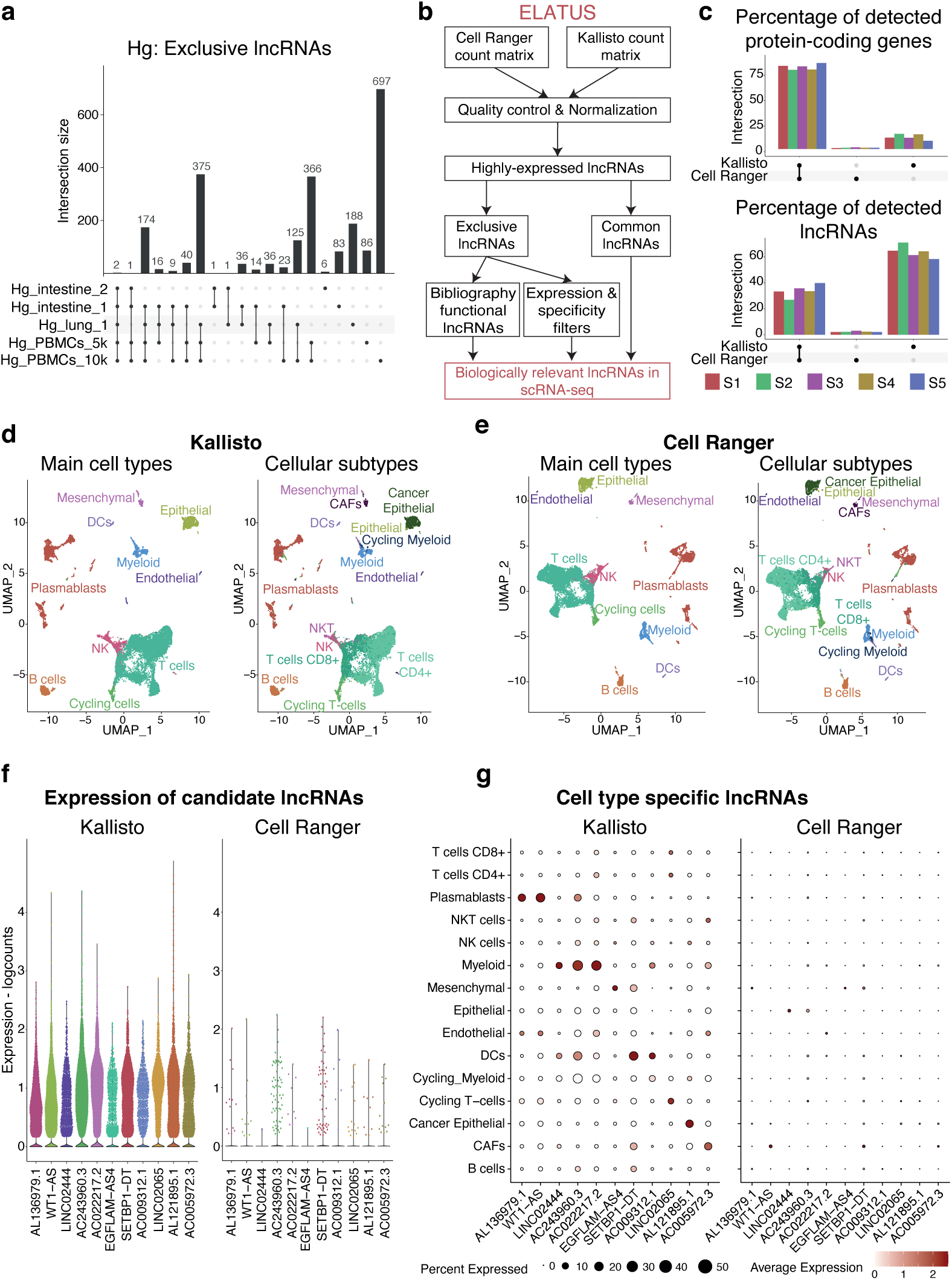
Biologically relevant lncRNAs are uncovered by ELATUS. **a,** UpSet plot displaying the overlap of lncRNAs exclusively found by Kallisto in the human scRNA-seq datasets analyzed. **b,** ELATUS workflow to uncover biologically important lncRNAs. It started by importing the raw count matrices obtained after preprocessing with both Cell Ranger and Kallisto. Next, there is a quality control step to distinguish empty droplets from cells, filtering of potential multiplets and cells with high mitochondrial content, followed by a normalization and clustering steps. Then, highly-expressed lncRNAs, both commonly detected by Cell Ranger and Kallisto and exclusively detected by Kallisto were selected. All the commonly detected lncRNAs were retained and from the exclusive lncRNAs, ELATUS retained those lncRNAs for which Cell Ranger assigned less than 10 counts, that were 40 times more expressed according to Kallisto than to Cell Ranger and that, according to Kallisto, had a SI > 0.15. ELATUS also retained the exclusive lncRNAs whose functionality has been independently validated by external studies. **c,** UpSet plot displaying, as a percetange, the overlap of: left) protein-coding genes, and right) lncRNAs detected by Kallisto and Cell Ranger in each sample. Only genes with more than 250 counts in more than 25 cells were considered in both panels. **d,** UMAP plots displaying the different TNBC cell population of: left) main cell types, and right) cell subtypes identified when preprocessing with Kallisto. **e,** UMAP plots displaying the different TNBC cell population of: left) main cell types, and right) cell subtypes identified when preprocessing with Cell Ranger. **f,** Violin plot showing the expression of some lncRNAs when preprocessing with Kallisto and Cell Ranger. **g,** DotPlots showing, with Kallisto and Cell Ranger, the averaged normalized expression in each cellular subtype of these lncRNAs.

Then, to further explore the potential of the ELATUS workflow in identifying lncRNAs with significant biological roles, we decided to investigate their expression in cells from triple-negative breast cancer (TNBCs) tumors. These highly aggressive breast tumors comprise various cell types, although our understanding of their transcriptional identity is still incomplete. We obtained five patient-derived TNBC fresh tumor biopsies and performed scRNA-seq that was then processed by ELATUS using both Cell Ranger and Kallisto (see Methods). After removing low-quality cells and poorly-expressed genes, we observed an important fraction of highly-expressed exclusive lncRNAs (1037 in total) in every sample, whereas most protein-coding genes were commonly detected (**Fig. 4c**), confirming our previous observations.

Next, we integrated the five TNBC samples, and cells were classified into major and minor cell types using canonical markers and reference datasets^50^ (**Fig. 4d-e, Extended Data Fig. 8b-c, Supplementary Data Table 1**). Further, we observed some lncRNAs overlooked by Cell Ranger that had unique expression patterns (**Fig. 4f**). Among them, using ELATUS we identified several cell type-specific candidates, such as *WT1-AS* and *AL133679.1*, which are plasmablasts-specific lncRNAs, or *AC009312.1* that is enriched in dendritic cells (**Fig. 4g**). Interestingly, we also recognized *AL121895.1*, that was specific of breast cancer epithelial cells and could be a potential marker of these tumorigenic cells. These findings highlight the importance of ELATUS for uncovering cell type and cancer type-specific lncRNAs in scRNA-seq experiments.

### ELATUS-identified *AL121895.1* is a *cis*-repressor that participates in triple negative breast cancer progression

Once established that our proposed workflow is able to detect cell type-specific lncRNAs that were previously missed by de facto preprocessing options, we next aimed to experimentally validate their expression patterns in different cell types. To do that, we selected *AL121895.1* and *WT1-AS*, which according to ELATUS are specific to breast cancer epithelial cells and plasmablasts, respectively (**Fig. 5a, Extended Data Fig. 8d**). We independently analyzed their expression in MDA-MB-231, a human epithelial breast cancer cell line, and in KMS-12-BM, a plasma cell line of multiple myeloma that represents later stages of B-cell differentiation^51–53^, similar to plasmablasts. Experimental detection by RT-qPCR confirmed the expression patterns of these lncRNAs that were found by ELATUS, since *AL121895.1* was significantly enriched in breast cancer cells, while *WT1-AS* had a significantly higher expression in multiple myeloma cells (**Fig. 5b**).

**Figure 5.**
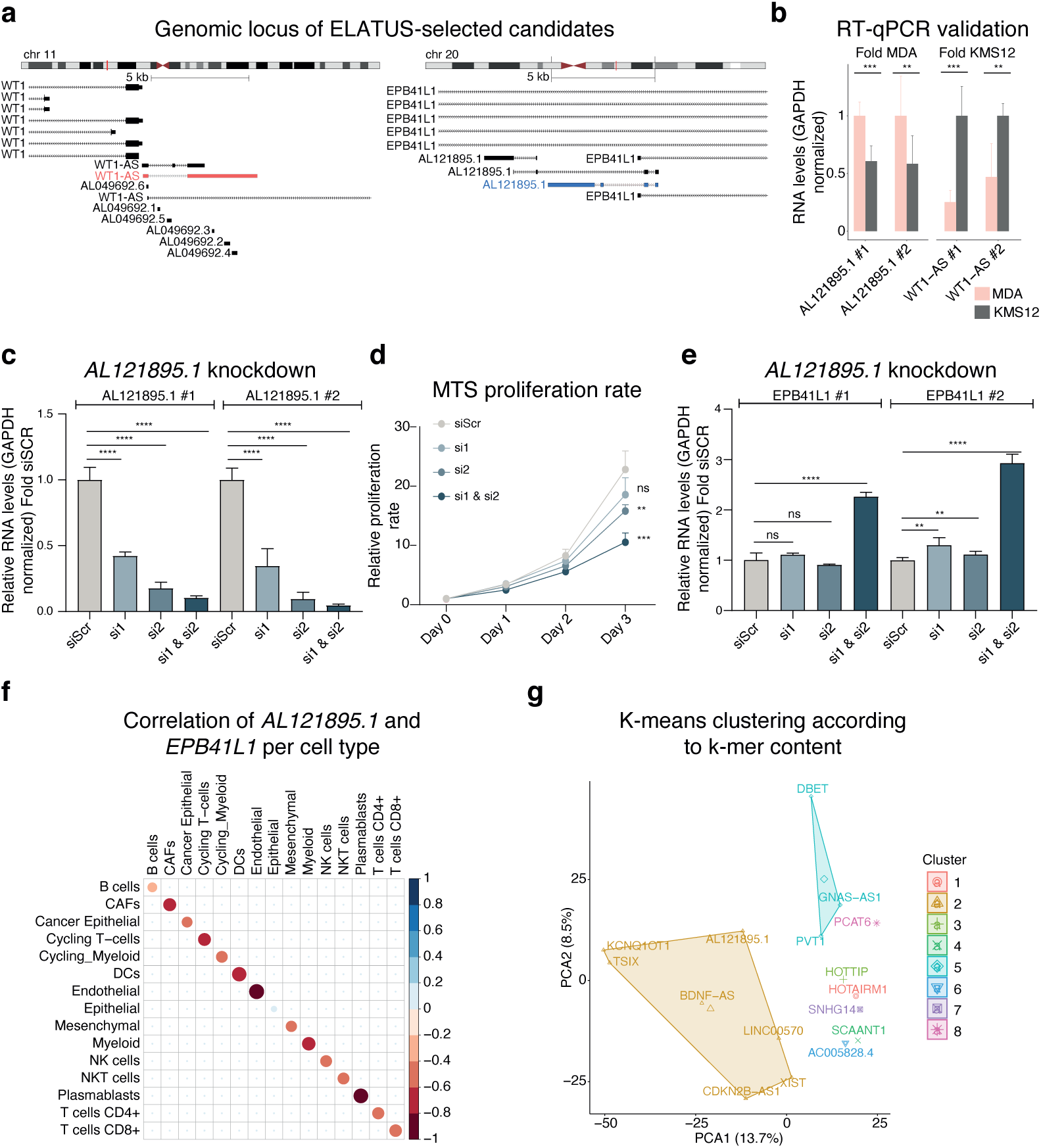
ELATUS-identified *AL121895.1* is a *cis*-repressor that participates in triple negative breast cancer progression. **a,** Genomic locus of left) *WT1-AS* and right) *AL121895.1*. In blue and red are represented the isoform of *WT1-AS* and *AL121895.1*, respectively, that contain most scRNA-seq reads assigned by Kallisto. **b,** RT-qPCR normalized RNA levels of *AL121895.1* and *WT1-AS* in MDA and KMS12 cell lines. *AL121895.1* and *WT1-AS* expression has been normalized with respect to MDA and KMS12, respectively. **c,** qPCR normalized RNA levels showing the expression of *AL121895.1* on MDA cells after treating them with scramble (siSCR) or knocked down with the siRNA1 (si1), siRNA2 (si2) and the combination of both siRNAs (si1 & si2). **d,** MTS proliferation assay of MDA cells measured during three days treating them with scramble (siSCR) or knocked down with the siRNA1 (si1), siRNA2 (si2) and the combination of both siRNAs (si1 & si2). **e,** RT-qPCR normalized RNA levels showing the expression of *EPB41L1* treating with the indicated siRNAs. Two-tailed student t-test for assessing statistical significance in the last four subpanels. **f,** Correlation plot of the normalized expression of *AL121895.1* and *EPB41L1* in each cellular subtype of the TNBC samples preprocessed with Kallisto. **g,** Functional classification by SEEKR using K-means clustering to find communities according to k-mer content of *AL121895.1* together with previously describer lncRNAs *cis*-activators and lncRNAs *cis*-repressors.

Given that *AL121895.1* is specific to breast cancer epithelial cells, we next investigated its function in these tumoral cells by knocking it down using siRNAs (**Fig. 5c, Extended Data Fig. 8d**). Interestingly, we observed that the knockdown of *AL121895.1* significantly decreased the proliferation rate of breast cancer cells (**Fig. 5d**), indicating a role in these tumorigenic cells. Furthermore, we noticed that its knockdown increased the expression of the antisense protein-coding gene, *EPB41L1* (**Fig. 5e, Extended Data Fig. 8e**), suggesting a previously undocumented *cis*-repressor function for *AL121895.1*. Inspection of the expression of *AL121895.1* and *EPB41L1* in the scRNA-seq of the TNBC samples, also confirmed a strong anticorrelation between them in each cell type (**Fig. 5f**). Further, since lncRNAs with similar k-mer content are functionally related^49^, we clustered, by K-means, the k-mer content of *AL121895.1* together with the k-mer composition of already described lncRNAs that are either *cis*-activators or *cis*-repressors^49^. The clustering grouped *AL121895.1* together with six other lncRNAs, from which five were proved *cis*-repressors (**Fig. 5g**), supporting its repressive role on its neighboring gene.

Together, these results demonstrate the biological significance of lncRNAs missed by Cell Ranger and highlight the need for ELATUS, the presented optimized scRNA-seq workflow in order to unlock cellular features encoded by lncRNAs.

## Discussion

It has been widely established that lncRNAs regulate gene expression and stability by different mechanisms at multiple levels^5,6,11^. However, given their high cell type specificity^7,8,10^, the current knowledge of lncRNAs could be greatly expanded with the application of single-cell technologies. Indeed, prior studies have showed that lncRNAs alone can identify cell types^1,54^ and that there are certain cell groups that can only be distinguished by them^3^. Nevertheless, compared to protein-coding genes, the annotation of lncRNAs is less well established and more prone to inaccuracies^15^, which complicates its correct quantification. In fact, as previously probed in bulk RNA-seq experiments^39^, pseudoaligners are promising alternatives to the closed-source and aligner-based Cell Ranger, which is the standard scRNA-seq preprocessing pipeline. In this work we have shown, in a wide and diverse set of public scRNA-seq datasets, that the detection and quantification of lncRNAs is severely affected by the preprocessing choice and the presented workflow tailored to lncRNAs, ELATUS, is essential, not only for defining an exhaustive collection of functional lncRNAs, but also for uncovering biologically important lncRNAs previously undetected.

Although previous studies have benchmarked the different preprocessing pipelines focusing on distinct metrics^27,32,35,36,42^ and noticed that Kallisto is able to detect a higher number of lncRNAs, they did not delve into the potential origin of this behavior. They either noted that it provided a comparable increment in the detection of both lncRNAs and protein-coding genes^36^, or attributed it to poorly expressed lncRNAs solely captured with Kallisto that did not provide a significant biological signal gain^27^. We hypothesize that this increased detection is not happening with Salmon due to its selective alignment strategy, in which the authors recommend including a set of decoy sequences to provide a more precise mapping^55^. Further, preceding investigations have alerted that there is a reduction in quantification accuracy of pseudoaligners due to the spurious mapping of intronic reads^29,42^, which can lead to false expression of particular genes^36^. The developers of Kallisto noted this phenomenon to be plausible but rare^32^. We provide multiple evidence supporting the validity of the proposed workflow for the detection of lncRNAs.

Due to the notably greater number of lncRNAs exclusively detected by Kallisto, we aimed to confirm whether they exhibit characteristics consistent with bona fide lncRNAs. To achieve this, we conducted a thorough characterization of the exclusive lncRNAs in comparison to those commonly captured. Our analysis suggests that the explanation attributing spurious alignments to poor quality mapping of low-complexity sequences does not fully apply to lncRNAs. Both common and exclusive lncRNAs are comparable in multiple parameters, and there is not a particular feature that causes the extra-detection. However, the fact that exclusive lncRNAs have a significantly higher number of exons supports the application of pseudoaligners and provides an alternative to the spurious mapping that also explains its increased detection rate. In order to mitigate the risk of including false transcript due to spurious mapping, here we implement ELATUS, which enriches for lncRNAs with functional features.

To determine the validity of ELATUS, the different expression patterns were tested computationally and experimentally. The multiple analyses performed suggested that the increased detection of lncRNAs is, at least in part, due to the identification of readily expressed functional transcripts^56^. Indeed, the expression of lncRNAs computed by Kallisto correlated more closely with ATAC-seq data than that of those detected by Cell Ranger. This indicates that pseudolignment effectively identified bona fide transcripts generated from regions of open chromatin. Moreover, the proposed computational workflow, ELATUS (available online as an R package), in addition to preserving robust and commonly detected lncRNAs, it unveils highly cell-type specific and biologically relevant lncRNAs from among the thousands of exclusive candidates missed by Cell Ranger. It is of particular relevance the previously uncharacterized lncRNA *AL121895.1*, specific of breast cancer epithelial cells and whose functionality at single-cell level could not be determined by standard preprocessing. Indeed, *AL121895.1* acts as a *cis*-repressor lncRNAs regulating *EPB41L1* expression and promoting TNBC progression. *EPB41L1* encodes a multifunctional protein that mediates interactions between the erythrocyte cytoskeleton and the overlying plasma membrane, although it is also expressed in other tissues^57,58^. Our results indicate that the regulation of *EPB41L1* by *AL121895.1* could represent a feature of breast tumor cells and a possible cancer vulnerability that could only be uncovered by ELATUS. The presented data evidences the potential impact of ELATUS to unveil important biological roles of lncRNAs and to expand the map of interactions, in individual cell populations, between the expression of a previously undetected by Cell Ranger lncRNA and the nearby protein-coding gene. Moreover, it exemplifies how scRNA-seq can inform outstanding mechanistic questions, such as the *cis* vs *trans* regulatory roles of lncRNAs.

It should be noted that the library preparation method could also influence the detection of lncRNAs. Conventional 10x Genomics scRNA-seq library preparation protocols target only polyA transcripts. Since an important fraction of lncRNAs are not polyadenylated^6^, these technologies cannot achieve a complete map of the transcriptome. Therefore, alternative scRNA-seq preparation protocols that capture both polyadenylated and non-polyadenylated transcripts, processed following the ELATUS workflow could potentially reveal an expanded number of functional non-coding transcripts that participate in important cellular functions.

Another direction that should be explored is the influence of the reference annotation. On the one hand, the application of an intronic reference, already recommended for snRNA-seq and that considers both mature and unmature RNAs has been widely discussed in the scRNA-seq community. However, the pre-mRNA reference poses a disjunctive when an exon of a gene is overlapped by an intron of another gene since every read falling in that region would be considered ambiguous (**Extended Data Fig. 10a-b**) without reaching a clear consensus on how to resolve these situations^59–61^. On the other hand, tissue-specific de novo annotation can greatly increase the detection of poorly annotated transcripts structures of specific lncRNAs^54^. Future investigation combining an improved reference annotation together with ELATUS could provide significant improvements in gene detection, especially for less studied biotypes such as lncRNAs.

Finally, with the proposed workflow, we favor the detection of lncRNAs with higher cell type specificity, as defined by the high specificity index (SI). We reason that this set of lncRNAs will include those with the most interesting biological features. However, while high cell specificity is recognized as a general characteristic of lncRNAs^7–10^, the existence of ubiquitous lncRNAs playing essential roles^62,63^ should not be excluded. Here, we propose an optimized computational workflow for analyzing scRNA-seq experiments that has the potential to unlock novel cellular features and transcriptional complexity, increasing the insights into cell identity and lncRNA biology.

## Methods

### Single-cell RNA-seq preprocessing pipelines

The following scRNA-seq preprocessing pipelines were benchmarked: Cell Ranger, STARsolo, Kallisto-Bustools (referred as Kallisto) and Salmon-Alevin (referred as Salmon). All pipelines were executed with the default recommended parameters in the user guides. For scRNA-seq analysis, Cell Ranger count was run in version 3.0.1, STAR in version 2.7.9, Kallisto in version 0.46.1, Bustools in version 0.40.0 and Salmon in version 1.4.0. For the pulmonary fibrosis dataset, since it was necessary to split cDNA sequence in more than one file, Kallisto was run in version 0.46.2 following authors indications^64^.

### Reference annotation and generation of the indexes

Human and mouse reference genome, transcriptome and annotation were downloaded from GENCODE^13^. Particularly, for human, hg38 (v37), and for mouse, mm10 (v27), were selected. We created the indexes for each preprocessing pipeline following recommended settings. Specifically, Cell Ranger and STARsolo were indexed against the entire genome, Kallisto against the transcriptome and Salmon was indexed to perform selective alignment to the transcriptome with full decoys as suggested by both the authors and independent benchmark studies^29,36,42,65^. The commands for preprocessing and generating the indexes for each preprocessing tool can be found in https://github.com/kikegoni/manuscript_scRNAseq_lncRNAs.

To analyze the evolution in the number of annotated protein-coding genes and lncRNAs, we analyzed the annotation provided by GENCODE since version 7, published in 2010, until the last version (v.44), published in December 2022.

### scRNA-seq public datasets

Different scRNA-seq prepared with 10x Genomics protocols, with both v2 and v3 chemistries, were used. Most of them were public, while sequencing data regarding TNBC samples were manually prepared. The description of each dataset as well as their link to access the sequencing data is provided in **Supplementary Data Table 4**.

### scRNA-seq quality control, gene detection and post-processing steps

Raw count matrices were used to standardize preprocessing pipelines as input for quality control, where we followed common scRNA-seq computational guidelines^66^. Specifically, emptydrops^67^ was applied to distinguish empty droplets from cells in each dataset processed with Cell Ranger, STARsolo and Kallisto. On the other side, Salmon, employs a whitelisting filtering strategy to filter empty droplets and it does not output the raw count data. To account for that and standardize filtering strategies, the minimum number of counts surviving emptydrops filtering in Cell Ranger, STARsolo and Kallisto was selected as an additional threshold to filter cells in Salmon with fewer counts than this defined threshold. Potential doublets were then identified and removed with scDblFinder^68^. Finally, cells with high mitochondrial content and an abnormally high number of counts were also filtered.

Once low-quality cells have been removed, in order to compare the detection and quantification of protein-coding genes and lncRNAs, poorly-expressed genes were also filtered by applying a gradient of thresholds on the expression. The different thresholds applied retained those genes with more than 1) 250 counts and present in more than 25 cells, 2) 100 counts and present in more than 10 cells, 3) 50 counts and present in more than 5 cells and 4) 25 counts and present in more than 3 cells. These thresholds were also applied in the characterization of the genes exclusively identified by Kallisto compared to genes commonly found ones. Due to the differences in the preserved number of cells in each scRNA-seq dataset, both absolute numbers and percentages have been used to represent the differences and the overlap of the detected highly-expressed genes by Kallisto and Cell Ranger.

After the quality control was completed, normalization was performed using logNormCounts function from scuttle R package^69^. Dimensionality reduction was conducted using runPCA, runTSNE and runUMAP functions from scater R package^70^. Next, clustering was performed on a generated shared nearest-neighbour (SNN)^71^ graph using the Louvain community detection algorithm to cluster the cells^72,73^. These clusters were manually annotated to cell types using canonical markers (**Supplementary Data Table 1**). Subtypes in TNBC integrated data were distinguished using the annotation program JIND^74^ selecting as a reference a dataset consisting of ∼45000 cells from TNBC samples^50^. We assessed the correlation (Spearman) between the normalized expression of *AL121895.1* and *EPBL41L1* per these cellular subtypes in each cell that expressed either *AL121905.1* or *EPBL41L1* (or both).

### Single-cell multiome analysis

In 3k PBMCs sequenced with single cell multiome, scATAC-seq raw data has been directly downloaded from 10X Genomics website^48^ on which scATAC-seq specific thresholds were first applied to remove low-quality nuclei. In particular, nuclei with very few or excessive ATAC-seq counts were filtered, as well as those with high nucleosome signal or little enrichment at the TSS^75^.

Gene expression data was obtained with Cell Ranger count (version 5.0.1) using --include-introns option. Regarding Kallisto (version 0.46.1) the index was generated to include both introns and exons using --workflow lamanno parameter.

For the raw RNA-seq matrices, those nuclei the fit the scATAC-seq quality control thresholds in both Cell Ranger and Kallisto were retained. Nuclei with very few RNA-seq counts or very high mitochondrial content were further removed. Next, ATAC-seq data from these high-quality nuclei were normalized using a Latent Semantic Indexing approach. “Weighted nearest neighbour” (WNN) analysis was then performed to integrate the ATAC-seq with the gene expression obtained by Cell Ranger and Kallisto. This integrated data was used for generating the clusters (following indications in scRNA-seq post-processing section) that were manually assigned to different cell types according to the expression of established marker genes.

To compare the similarity between the ATAC-seq signal and the RNA-seq gene expression, the GeneActivity function from Signac R package^76^ was applied to obtain a gene activity matrix by counting the scATAC-seq fragments that fall in each gene body (+2Kb upstream from the TSS). Then, for every high-quality nucleus we compared (student’s t-test) the number of genes that have simultaneous ATAC-seq-signal and RNA-seq expression when preprocessed with Cell Ranger or with Kallisto. A gene was defined to be simultaneously activated if its ATAC-seq signal and its RNA-seq expression were higher than a gradient of defined thresholds. Specifically, from less to more restrictive thresholds; if they had at least 1) 1 read in ATAC-seq and 1 UMI in RNA-seq, 2) 3 reads in ATAC-seq and 3 UMIs in RNA-seq, 3) 6 reads in ATAC-seq and 6 UMIs in RNA-seq, 4) 6 reads in ATAC-seq and 11 UMIs in RNA-seq and 5) 11 reads in ATAC-seq and 11 UMIs in RNA-seq.

The differences, per cell, were also represented as an odds ratio (in log2 scale) showing the likelihood of having more genes simultaneously activated with Kallisto than with Cell Ranger. Further, the ratio (in log2 scale), of the number of nuclei for which there were more simultaneous activation when the snRNA-seq was processed with Kallisto than with Cell Ranger has been computed. A positive ratio indicates a better correspondence between ATAC-seq and Kallisto than between ATAC-seq and Cell Ranger, while a negative ratio indicates the opposite.

### Exclusive vs common genes: Length, number of exons, repeat content and k-mer analysis

To investigate the length, the number of exons and repeat content of both exclusive and commonly detected genes in every scRNA-seq dataset, the longest isoform of each gene was selected. The annotation of repeats for both human and mouse genomes was downloaded from RepeatMasker (version 4.1.5)^77^, where we considered all distinct types of repeats with the exception of microsatellites repeats. We calculated the ratio between the percentage of the sequence of exclusive features covered by repeats divided by the percentage of the sequence of common features covered by repeats.

We compared the k-mer content of all lncRNAs using SEEKR^49^, a software developed for sequence evaluation through k-mer content, where we calculated the 6-mer functionally-related communities using the canonical isoform of each gene following the default recommended settings by the authors^78^. For the K-means (stats R package) clustering of *AL121895.1* according its k-mer content and the k-mer content of previously described cis-activators or cis-repressors^49^, we considered the isoform *AL121895.1-EST00000441208*. This is the isoform to which Kallisto assigned most of scRNA-seq reads and for which qPCR primers and siRNAs were designed.

### Exclusive vs common genes: Specificity index

In order to implement the Specificity Index (SI), in line with other methods^4,54^, each scRNA-seq datasets was clustered across distinct ranges of number and sizes of clusters, from fewer and bigger clusters to more but smaller clusters. We started by splitting the cells in 5-9 clusters, then 10-15 clusters, then 16-22 clusters and finally generating a very detailed subdivision with at least 23 clusters. The SI metric was then designed in order to inform if a lncRNA is more specific of big or small subpopulations. We implemented the SI following the Shannon-Entropy specificity (HS) formulation defined in TSPEX, a library with several specificity metrics^79^. So, in order to define the SI for each gene *g*, we first calculated its mean expression *x_i_* in each cluster, *i* = 1,2…*n*, where *n* is the number of clusters. Next, we calculated for each gene, the proportion of mean expression in each cluster, *P_i_*:

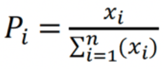

Finally, using the entropy HS formulation, we implemented the SI metric, where we assessed if each gene is expressed in fewer and localized clusters or if its expression is more broadly expressed. Concretely the SI, for each gene, was defined as:

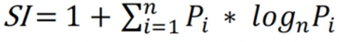

The SI is ranked from [0-1], from genes very ubiquitously expressed to very cluster-specific genes. A gene whose expression is equally distributed across different clusters will have a SI of 0, while if a gene is exclusively expressed in one cluster its SI will be 1.

### ELATUS workflow defines a collection of functional lncRNAs

On the one side, to get the list of biologically relevant lncRNAs, we analyzed the 288 CRISPR functionally validated lncRNAs in multiple human cell lines^10^. Using a hypergeometric test, we tested the significance of their overlap with those lncRNAs exclusively identified by Kallisto, in every human scRNA-seq. In total, there was an overlap of 173 Kallisto-exclusive lncRNAs.

On the other side, we implemented ELATUS in order to capture the highly-expressed lncRNAs commonly detected and to inspect the highly-expressed lncRNAs exclusively identified with Kallisto to assess their biological relevance. Therefore, we started by importing the raw count matrices obtained with both Cell Ranger and Kallisto and we performed emptydropts to distinguish empty droplets from cells. Next, we removed potential multiplets and performed a quality-control filtering, followed by a normalization and clustering steps. Further, we integrated samples from the same tissue. Then, highly-expressed lncRNAs (i.e. those with more than 250 counts and present in more than 25 cells), both commonly detected by Cell Ranger and Kallisto and exclusively identified by Kallisto were selected. All the commonly detected lncRNAs were retained, whereas from the exclusive lncRNAs ELATUS retained those lncRNAs for which Cell Ranger assigned lass than 10 counts, that were 40 times more expressed according to Kallisto than to Cell Ranger and that, according to Kallisto, had an SI > 0.15. ELATUS also retained the exclusive lncRNAs whose functionality has been independently validated by external studies. To include a representative set of cluster sizes to calculate the SI, scRNA-seq datasets were divided in different cluster sizes, from 10 to 19 clusters (**Supplementary Data Table 4**). ELATUS, which is openly available as an R package in https://github.com/kikegoni/ELATUS, has been executed with these restrictive thresholds to ensure biological relevance and to minimize the risk of false expression caused by spurious mapping.

### Statistical analysis and data plotting

Post-processing analysis were performed in R (version 4.1.2). Barplots and violin plots were represented with ggplot2 (v.3.4.2), where ggpubr (v.0.4.0) was used to test statistical test significance. The specific statistical test for each analysis is detailed in its figure caption, where *ns* represents p-value > 0.1, *\** represents p-value <= 0.1, *\*\** represents p-value <= 0.05, *\*\*\** represents p-value <= 0.005 and *\*\*\*\** represents p-value <= 0.0005.

From Seurat (v.4.0.1)^80^, DimPlot function was used to plot UMAP dimensionality reduction plots and DotPlot and FeaturePlots functions were applied to evaluate gene expression in different cell types and dimensionality reduction spaces, respectively. UpSet plots were generated with UpSetR (v.1.4.0)^81^ and ggupset (v.0.3.0). In the analysis of single-cell multiome data, Signac R package (v. 1.9.0)^76^ was applied to create the coverage and expression plots.

### Cell lines and growth conditions

MDA-MB-231 cells were cultured in DMEM (GIBCO), supplemented with 10% fetal bovine serum (GIBCO) and 1x penicillin/streptomycin (Lonza), while KMS-12 cells were grown in RPMI-1640 (GIBCO) medium with 20% fetal bovine serum (GIBCO), 1% penicillin, and 2% Hepes. All of them were maintained at 37°C and 5% CO_2_.

### RNAi

For RNA knockdown, siRNAs, which were designed using the i-Score designer tool and purchased from Sigma (**Table S1**), were transfected for 24 hours at 40 nM final concentration. MDA-MB-231 cells were transfected with Lipofectamine 2000 (Invitrogen) in Serum-free Opti-MEM (GIBCO), following manufacturer instructions.

**Table S1.**
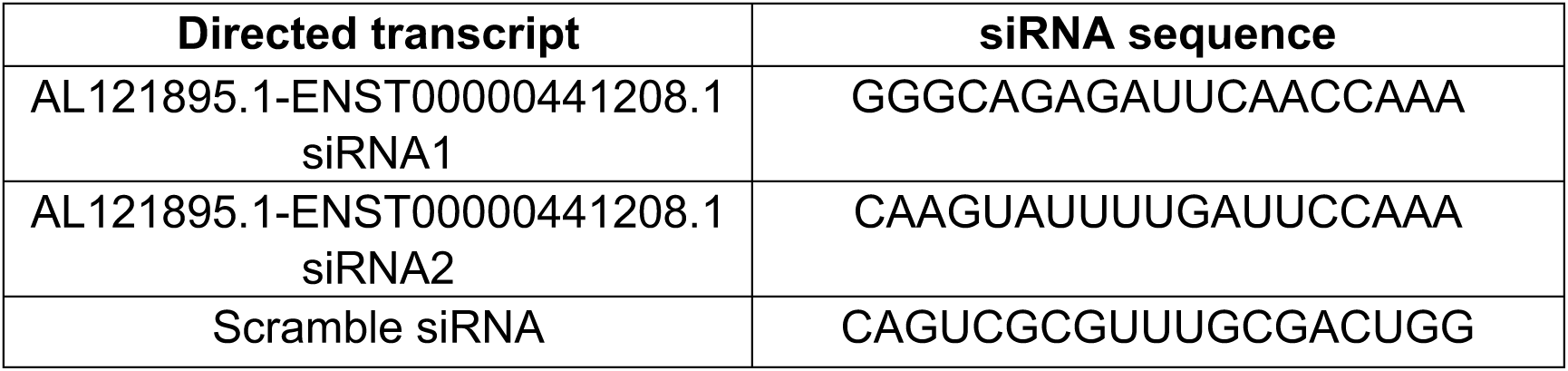
List and sequence of siRNAs designed to knockdown lncRNAs.

### Proliferation assay

Cell proliferation was measured using the CellTiter96 Aqueous Non-Radioactive Cell Proliferation Assay (MTS) kit (Promega). After 24 hours transfection, 1000 MDA cells were cultured in M-96 plate wells. Every 24 hours, 20 µL of MTS reagent (Promega) were added to culture media, and incubated for 2 hours, prior to 490 λ measurement. Triplicate measures were normalized to day 0, and statistical differences between control and experimental conditions at day 3 were calculated with a two-tailed student t-test.

### RNA extraction, processing, and RT-qPCR

Cell preparations were fixed with TRIzol (Sigma), and RNA precipitated with isopropanol. RNA extraction was followed by Turbo DNAse (Invitrogen) digestion for 30 minutes at 37°C. For RT-qPCR, 1 µg RNA was reverse-transcribed using the High-Capacity cDNA Reverse Transcription Kit (Applied 30 Biosystem) with random hexamer primers, following manufacturer instructions. The obtained cDNA was analysed by quantitative PCR (qPCR) using iTaq Universal SYBR Green supermix (Bio-Rad) in a ViiA™ 7 Real-Time PCR System machine (ThermoFisher), all reactions performed in quadruplicate. GAPDH RNA levels were used for normalization. Statistical differences between relative RNA levels were calculated by unpaired two-tailed Student’s *t*-test. RT-qPCR primers were self-designed or designed with the NCBI Primer designing tool, and purchased from Metabion (**Table S2**).

**Table S2.**
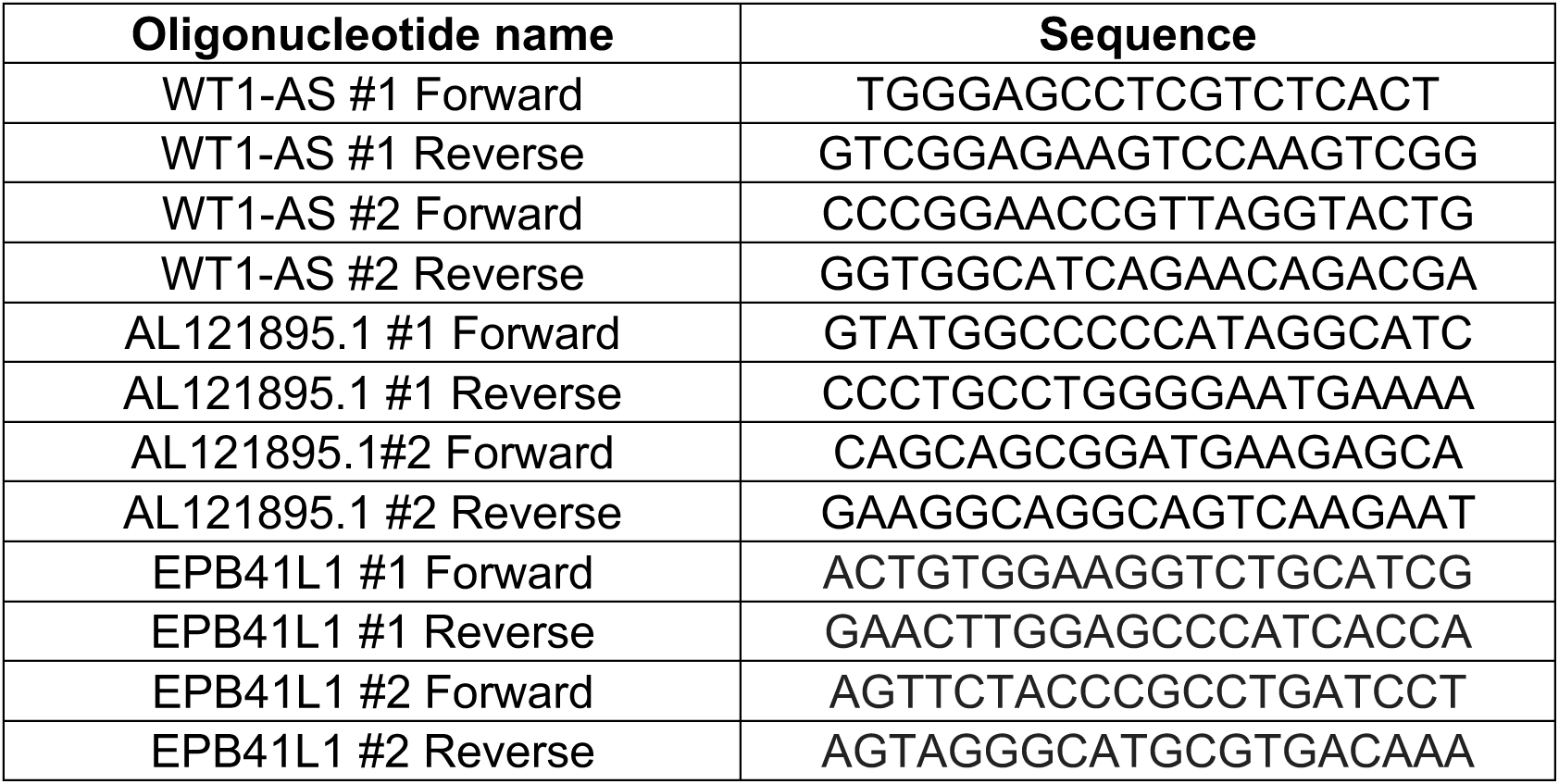
List and sequence of oligonucleotides designed for RT-qPCR.

### Human samples

All patients participating in the study provided written informed consent. The study and the use of all clinical materials have been approved by the Research Ethics Committee of the University of Navarra under decision number 2021.058Mod1. Samples and data from patients included in the study were provided by the Biobank of the University of Navarra and were processed following standard operating procedures approved by the Ethical and Scientific Committees.

### TNBC sample preparation and preparation of the scRNA-seq libraries

Five biopsies of tissue from different patients of breast cancer were processed following manufacturer’s instructions of Human Tumor Dissociation Kit from Miltenyi Biotec. Briefly, 2-3 tissue cylinders per patient were cut into small slices of 2–4 mm, digested with enzymes H, R and A, and dissociated with gentleMACS Dissociator (Miltelnyi Biotec). After 30 minutes incubation at 37°C and a short centrifugation step (1800 rpm, 1 minute at 4°C), sample material at the bottom of the tube was collected. cell suspension was then moved to a Falcon cell strainer (70 μm) placed on a 50 mL tube, washed with 20 mL of DMEM, and centrifuged at 1800 rpm for 5 minutes at 4°C. Pelleted cells were washed with 1ml of PBS 1X 0.05%BSA, supplemented with 5ul RNAse OUT (Invitrogen), and transferred to a Dolphin tube (Sorenson) where cell suspension was centrifuged at 1800 rpm for 5 minutes at room temperature. Viability (>70%) of resuspended cells was corroborated with cellometer (Nexcelom).

The transcriptomes of 16,000-20,000 cells were examined using Single Cell 3’ Reagent Kits v3.1 (10X Genomics) according to the manufacturer’s instructions. Briefly, 17000 to 20000 cells were loaded at a concentration of 1000 cells/µL on a Chromium Controller instrument (10X Genomics) to capture single cells in gel bead-in-emulsions (GEMs). In this step, each cell was encapsulated with primers containing a fixed Illumina Read 1 sequence, a cell-identifying 16 bp 10x Genomics barcode, a 12 bp Unique Molecular Identifier (UMI) and a poly-dT sequence. Upon cell lysis, reverse transcription yielded full-length, barcoded cDNA. This cDNA was then released from the GEMs, PCR-amplified and purified with magnetic beads (SPRIselect, Beckman Coulter). Enzymatic Fragmentation and Size Selection was used to optimize cDNA size prior to library construction. Fragmented cDNA was then end-repaired, A-tailed and ligated to Illumina adaptors. A final PCR-amplification with barcoded primers allowed sample indexing. Library quality control and quantification was performed using Qubit 3.0 Fluorometer (Life Technologies) and Agilent’s 4200 TapeStation System (Agilent), respectively. Sequencing was performed in a NextSeq2000 (Illumina) (Read1: 28; Read2: 91; i7 index: 8) at an average depth of 500000 reads/sample.

### Data availability

TNBC scRNA-seq data has been publicly stored in NCBI with the following identifier: GSE246142.

### Code availability

The code and scripts to reproduce the results is freely available and can be accessed in https://github.com/kikegoni/manuscript_scRNAseq_lncRNAs. ELATUS R package for elucidating biologically relevant lncRNA annotated transcripts using scRNA-seq is available at https://github.com/kikegoni/ELATUS.

## Figures

**Extended Data Figure 1.**
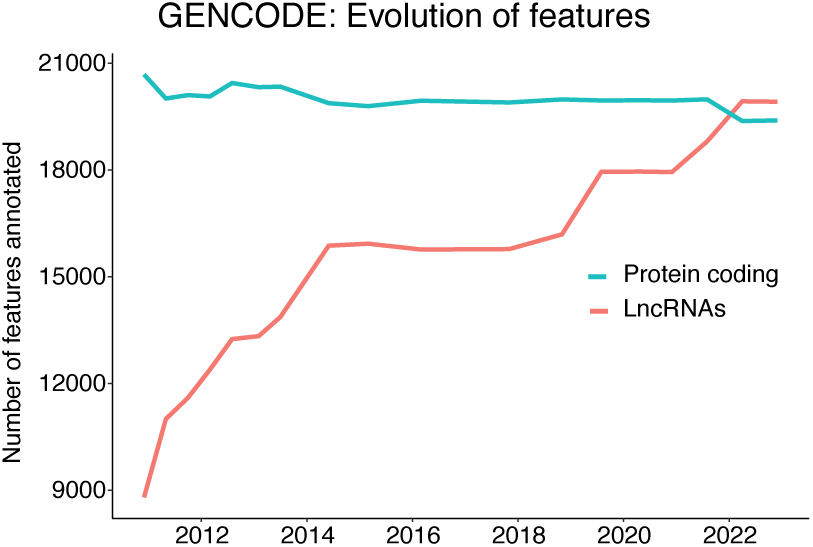
Evolution of the number of protein-coding genes and lncRNAs annotated by GENCODE from 2010 until 2022.

**Extended Data Figure 2.**
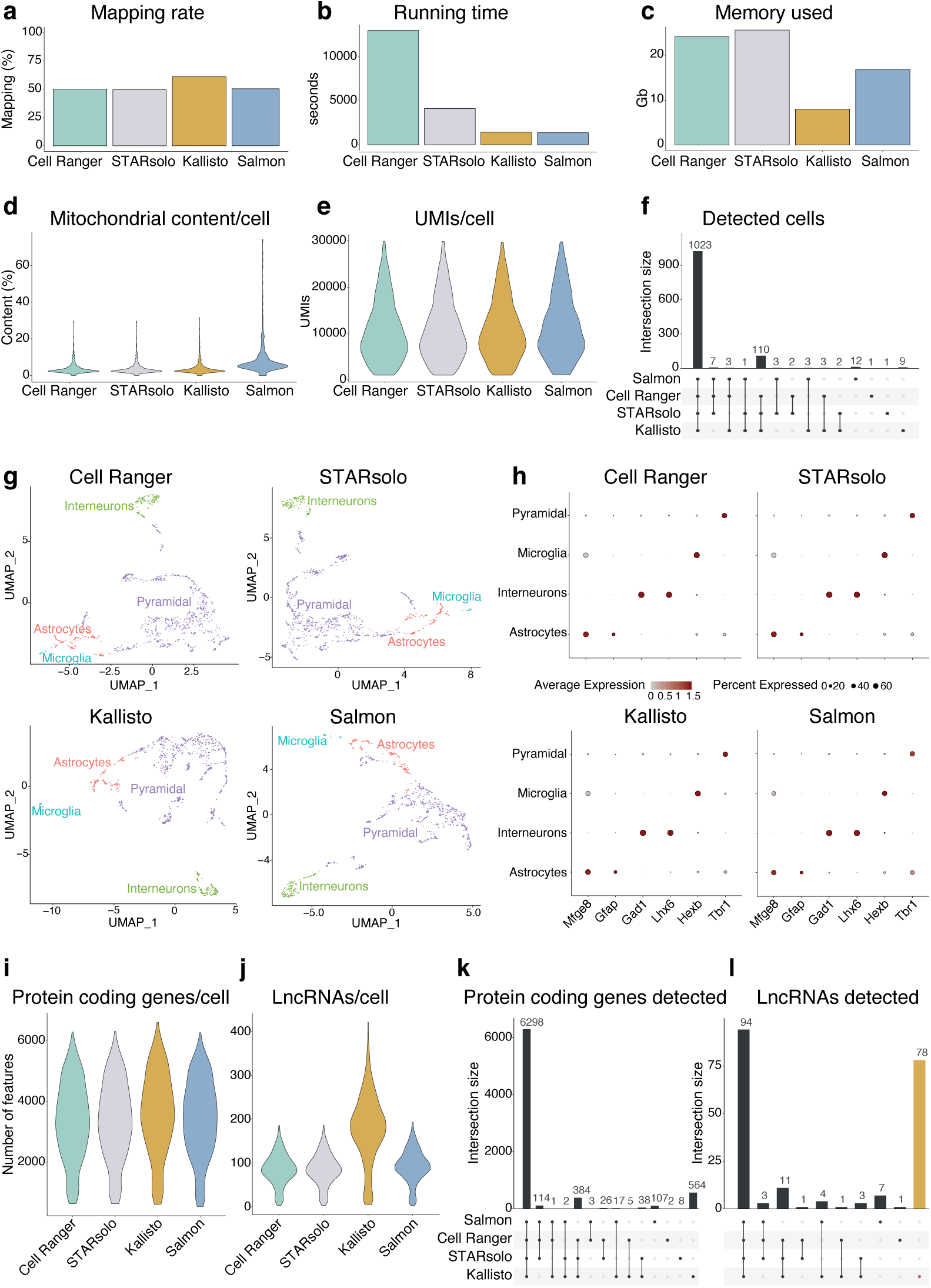
Preprocessing choices strongly affect lncRNA detection in a scRNA-seq dataset consisting of 1k brain cells from an E18 mouse. **a,** Mapping rate by each pipeline. **b,** Running time per pipeline. **c,** Maximum memory used per pipeline. **d,** Mitochondrial content per cell by each pipeline. **e,** Number of UMI counts per cell by each pipeline. **f,** UpSet plot showing the overlap of retained high-quality cells by each pipeline. **g,** UMAP plots displaying the populations of cell types identified across pipelines. **h,** DotPlots showing across different pipelines, the averaged normalized expression, in each cell type, of canonical marker genes. **i,** Number of detected protein-coding genes per cell across pipelines. **j,** Number of detected lncRNAs per cell across pipelines. **k,** UpSet plot displaying the overlap of highly-expressed protein-coding genes (more than 250 counts and present in more than 25 cells) per pipeline. **i,** UpSet plot displaying the overlap of highly-expressed lncRNAs (more than 250 counts and present in more than 25 cells) per pipeline.

**Extended Data Figure 3.**
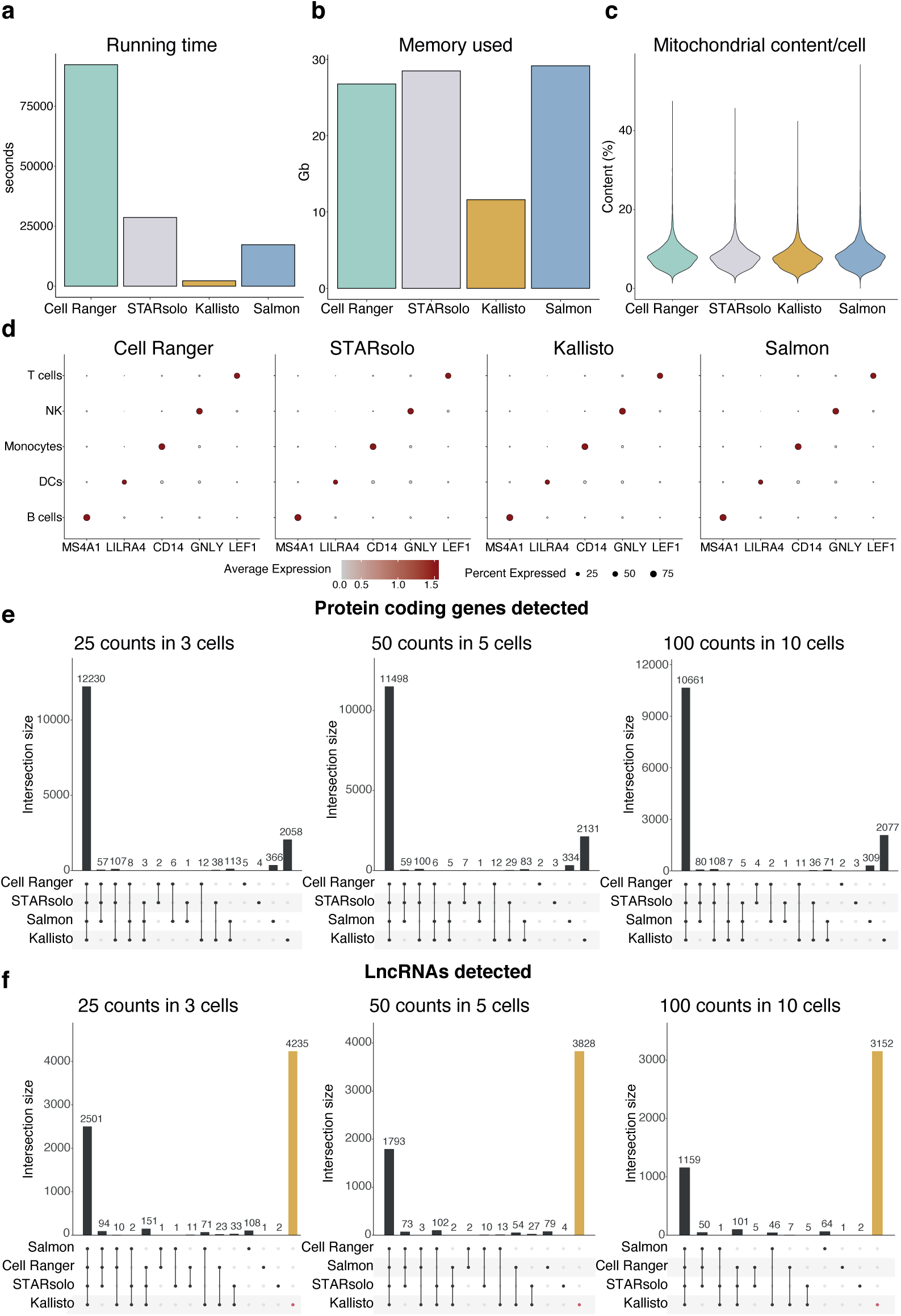
Preprocessing choices strongly affect lncRNA detection in a scRNA-seq dataset consisting of 10k human PBMCs from a healthy donor. **a,** Running time per pipeline. **b,** Maximum memory used per pipeline. **c,** Mitochondrial content per cell by each pipeline. **d,** DotPlots showing across different pipelines, the averaged normalized expression, in each cell type, of canonical marker genes. **e,** UpSet plot displaying the overlap of highly-expressed protein-coding genes across pipelines in different thresholds. Specifically, in left) more than 25 counts and present in more than 3 cells, middle) more than 50 counts and present in more than 5 cells, right) more than 100 counts and present in more than 10 cells. **f,** UpSet plot displaying the overlap of highly-expressed lncRNAs across pipelines in different thresholds. Specifically, in left) more than 25 counts and present in more than 3 cells, middle) more than 50 counts and present in more than 5 cells, right) more than 100 counts and present in more than 10 cells.

**Extended Data Figure 4.**
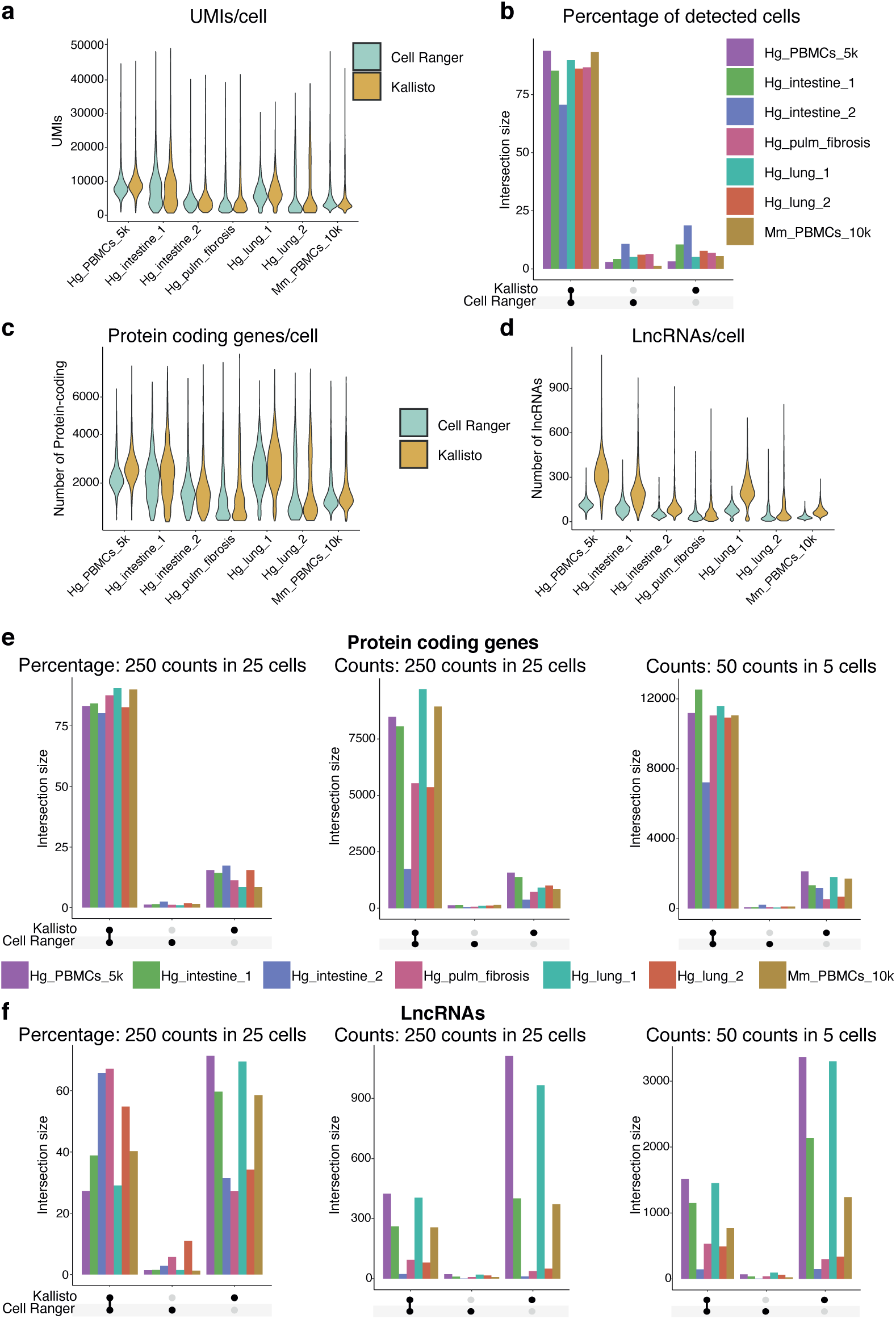
Preprocessing choices strongly affect lncRNA detection in an expanded and diverset set of scRNA-seq datasets. **a,** Number of UMI counts per cell obtained with Cell Ranger and Kallisto in each dataset. **b,** UpSet plot displaying the percentage of the detected high-cells which are individually or commonly retained by Cell Ranger and Kallisto across datasets. **c,** Number of detected protein-coding genes per cell by Cell Ranger and Kallisto in each dataset. **d,** Number of detected lncRNAs per cell by Cell Ranger and Kallisto in each dataset. **e,** UpSet plot displaying, for highly-expressed protein-coding genes, the left) percentage of them that have more than 250 counts in more than 25 cells that overlaps between Cell Ranger and Kallisto middle) overlap of them that have more 250 counts in more than 25 cells between Cell Ranger and Kallisto, right) overlap of them that have more 50 counts in more than 5 cells between Cell Ranger and Kallisto. **f,** UpSet plot displaying, for highly-expressed lncRNAs, the left) percentage of them that have more than 250 counts in more than 25 cells that overlaps between Cell Ranger and Kallisto middle) overlap of them that have more 250 counts in more than 25 cells between Cell Ranger and Kallisto, right) overlap of them that have more 50 counts in more than 5 cells between Cell Ranger and Kallisto.

**Extended Data Figure 5.**
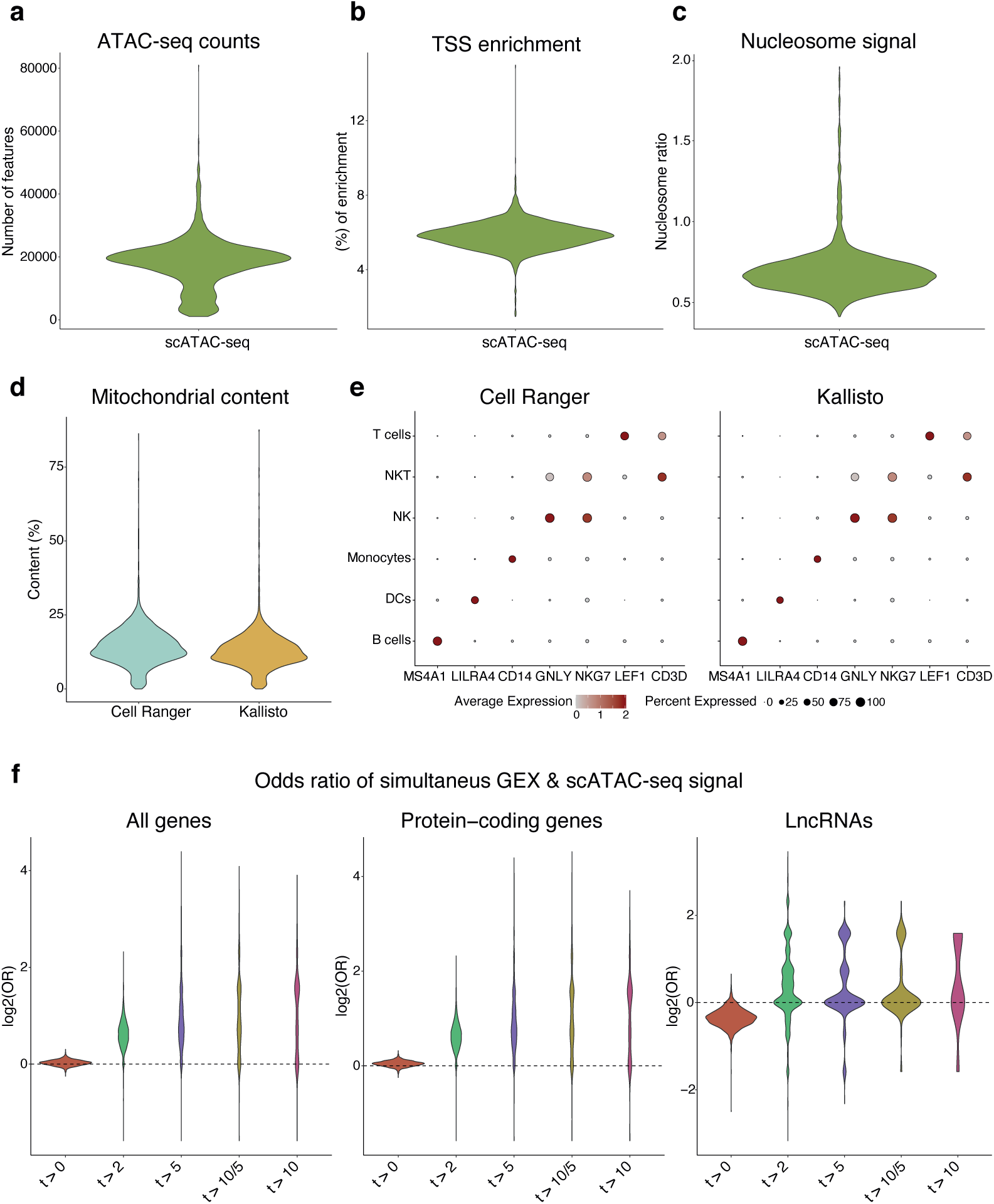
scATAC-seq multiome indicates the optimal preprocessing alternative for lncRNA quantification. **a,** ATAC-seq counts for each nucleus that fits ATAC-seq quality control thresholds. **b,** Transcription Start Site (TSS) enrichment for each nucleus that fits ATAC-seq quality control thresholds. **c,** Nucleosome signal for each nucleus that fits ATAC-seq quality control thresholds. **d,** Mitochondrial content per nuclei by Cell Ranger and Kallisto. **e,** DotPlots showing across different pipelines, the averaged normalized scRNA-seq expression, in each cell type, of canonical marker genes. **f,** Odds ratio displaying the probability of having more simultaneous activation with Kallisto than with Cell Ranger across different thresholds for defining a gene to be simultaneously activated. Specifically, the x-axis represents the different thresholds used for quantifiying only a gene as simultaneously activated if it had: (t>0) at least 1 UMI in RNA-seq and 1 read in ATAC-seq and, (t>2) at least 3 UMIs in RNA-seq and 3 reads in ATAC-seq, (t>5) at least 6 UMIs in RNA-seq and 6 reads in ATAC-seq, (t>10/5) at least 11 UMIs in RNA-seq and 6 reads in ATAC-seq and (t>10) at least 11 UMIs in RNA-seq and 11 reads in ATAC-seq. For each nucleus we have considered the expression of all genes (left), only protein-coding genes (middle) and only lncRNAs (right).

**Extended Data Figure 6.**
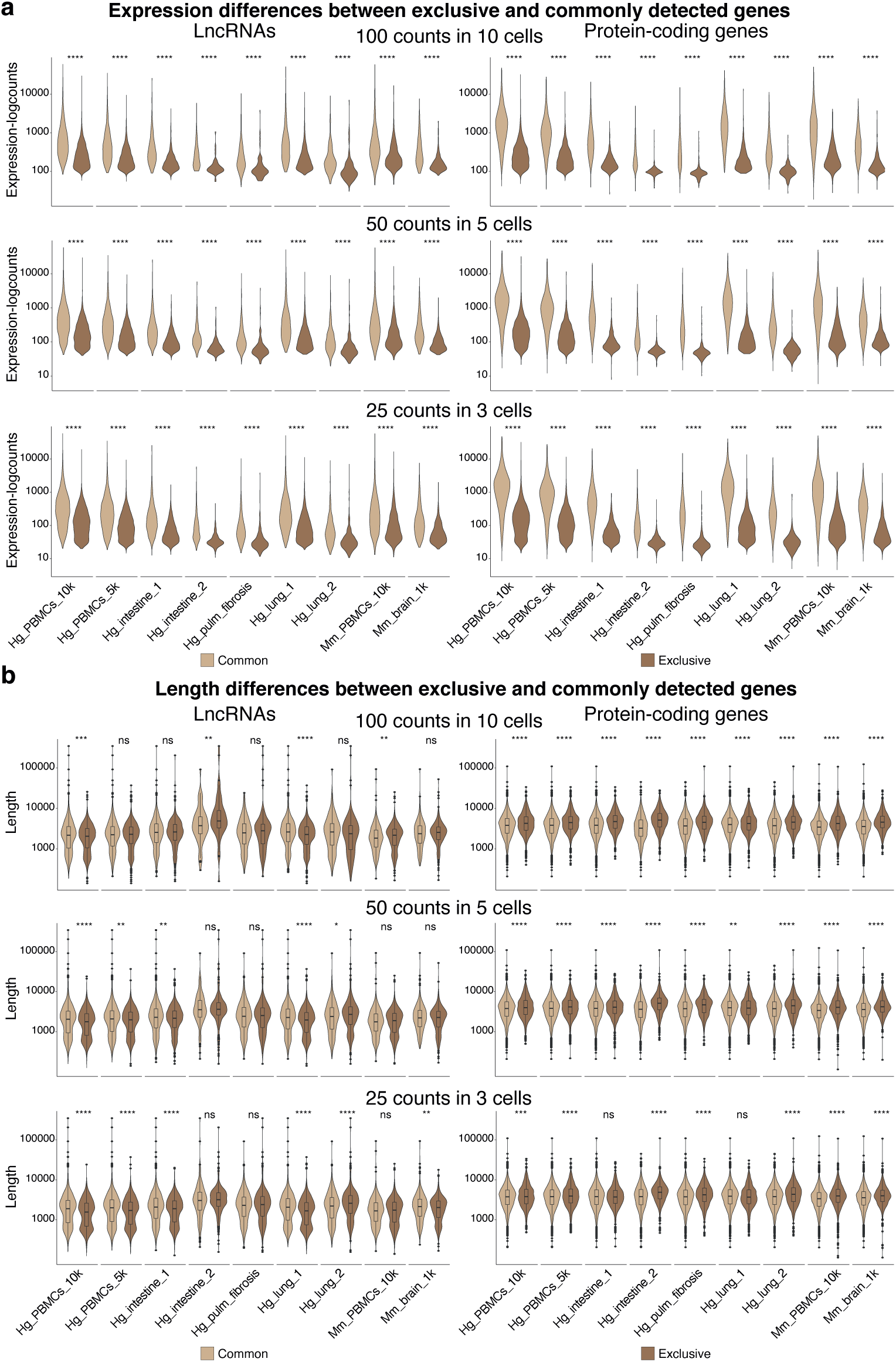
Exclusive and common lncRNAs have similar expression and length distributions. **a,** Normalized expression differences and **b,** Length differences of (left) lncRNAs exclusively found by Kallisto compared to lncRNAs commonly identified by all pipelines and (right) protein-coding genes exclusively found by Kallisto compared protein-coding genes commonly identified by all pipelines. Significance was assessed with a two-tailed Wilcoxon test in (**a**) and (**b**). Thresholds in (**a**) and (**b**) applied for removing poorly-expressed genes were; up) only genes with more than 100 counts and present in more than 10 cells were considered, middle) only genes with more than 50 counts and present in more than 5 cells were considered, down) only genes with more than 25 counts and present in more than 3 cells were considered.

**Extended Data Figure 7.**
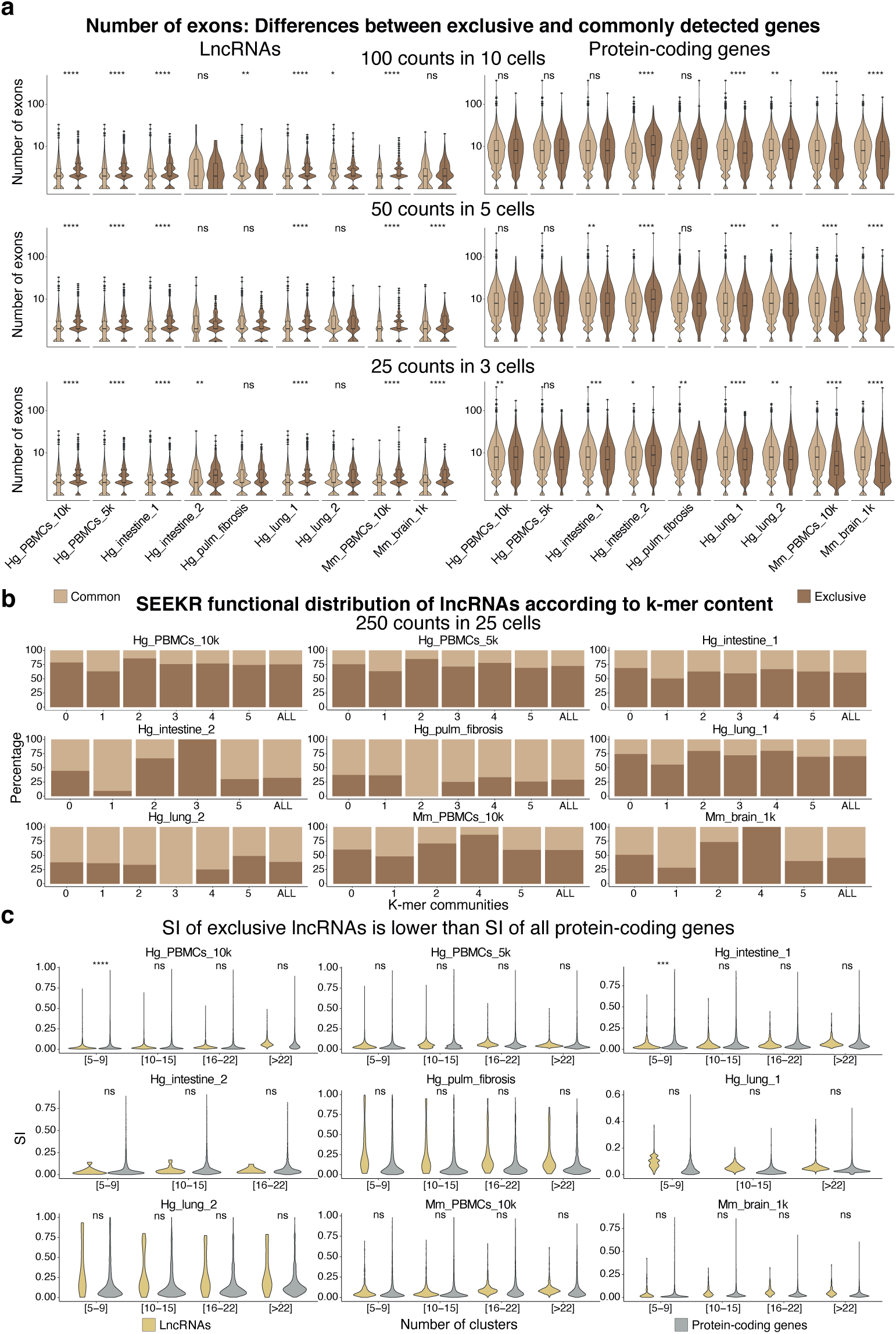
Exclusive lncRNAs are highly specific and have a higher number of exons than those commonly detected. **a,** Differences in the number of exons of (left) lncRNAs exclusively found by Kallisto compared to lncRNAs commonly identified by all pipelines and (right) protein-coding genes exclusively found by Kallisto compared to protein-coding genes commonly identified by all pipelines. Significance was assessed with a two-tailed Wilcoxon test. Thresholds applied for removing poorly-expressed genes were; up) only genes with more than 100 counts and present in more than 10 cells were considered, middle) only genes with more than 50 counts and present in more than 5 cells were considered, down) only genes with more than 25 counts and present in more than 3 cells were considered. **b,** Percentage of exclusive and common lncRNAs across six SEEKR functional communities [0-5] according to k-mer content of lncRNAs. The percentage of exclusive and common lncRNAs in total also included to evaluate differences (column *ALL*). To remove poorly-expressed genes we considered only genes with more than 250 counts and present in more than 25 cells were considered. **c,** Specificity index (SI) differences to test if the SI of lncRNAs exclusively identified by Kallisto is significantly lower (one-tailed Wilcoxon test) than the SI of protein-coding genes. The SI distributions were calculated across distinct ranges of number and sizes of clusters, from fewer and bigger to more and smaller clusters. Starting with 5-9 clusters, then 10-15 clusters, then 16-22 clusters and finally generating a specially detailed subdivision with at least 23 clusters. To remove poorly-expressed genes we considered only genes with more than 250 counts and present in more than 25 cells were considered.

**Extended Data Figure 8.**
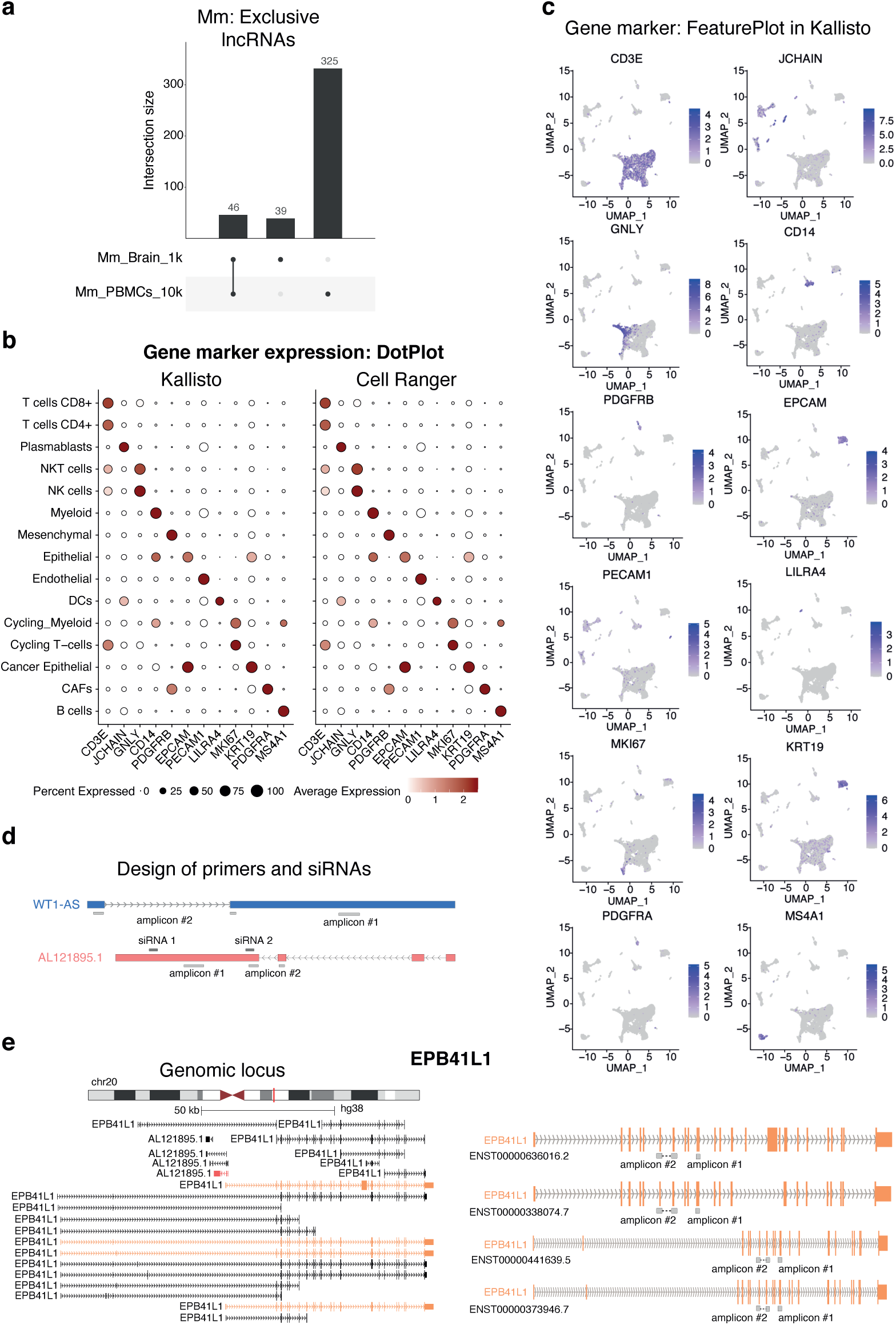
Functional lncRNAs are uncovered by ELATUS. **a,** UpSet plot displaying the overlap of lncRNAs exclusively found by Kallisto in the mouse scRNA-seq datasets analyzed. **b,** DotPlots showing the averaged normalized expression, in each cellular subtype, of canonical marker genes when preprocessing with Kallisto and Cell Ranger **c,** UMAPs showing the normalized expression of canonical marker genes when preprocessing with Kallisto **d,** Design of the set of qPCR primers and siRNAs used for *WT1-AS* and *AL121895.1*. In blue and red are represented the isoform of *WT1-AS* and *AL121895.1*, respectively, that contain most scRNA-seq reads assigned by Kallisto **e,** Left) Genomic locus of the protein-coding gene *EPB41L1* and right) Design of the pair of qPCR primers used for the isoform containing most scRNA-seq reads assigned by Kallisto.

**Extended Data Figure 9.**
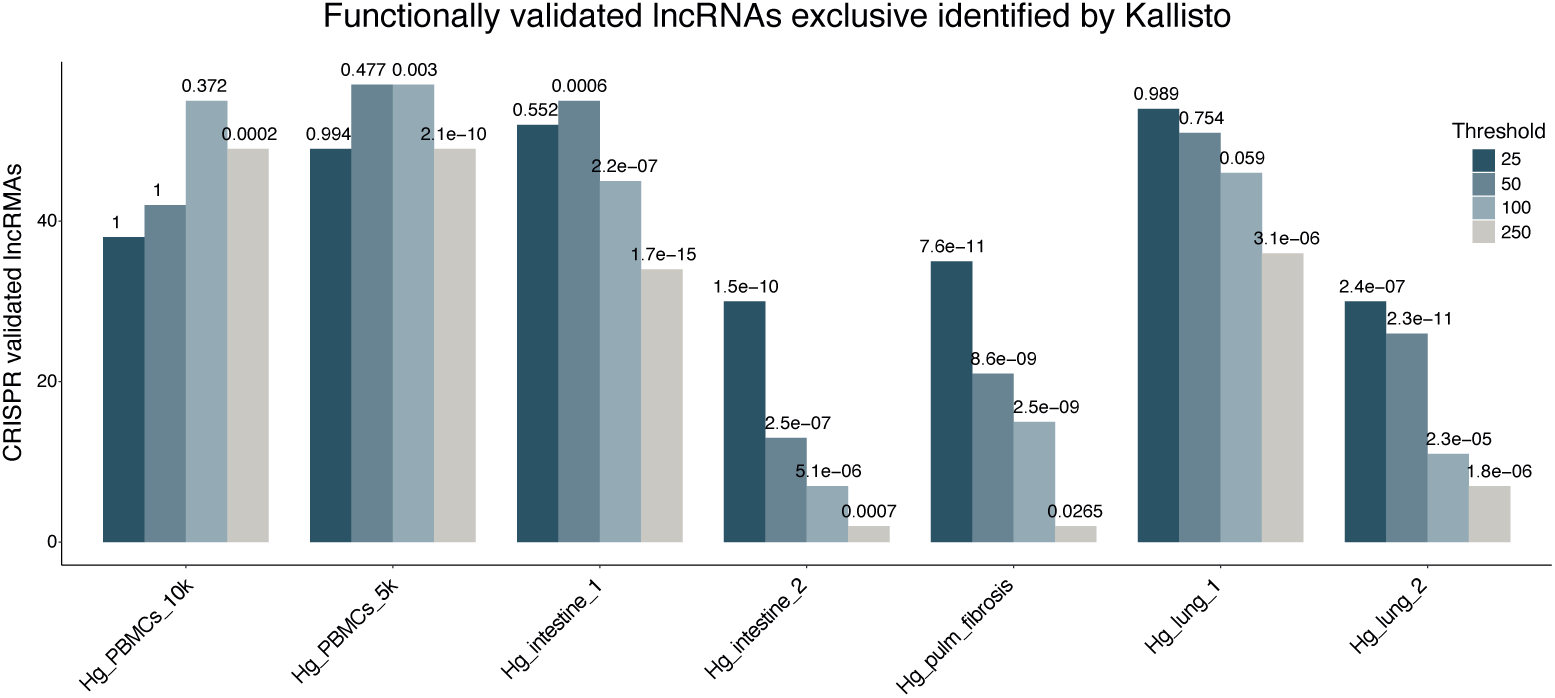
Exclusive lncRNAs are significantly enriched among CRISPR functionally validated lncRNAs. Hypergeometric test performed to test the significance of the overlap. Different thresholds on expression for removing lowly-expressed genes were considered, from left to right; more than 25 counts and present in more than 3 cells (dark blue), more than 50 counts and present in more than 5 cells (blue), more than 100 counts and present in more than 10 (light blue) cells and more than 250 counts and present in more than 25 cells (grey).

**Extended Data Figure 10.**
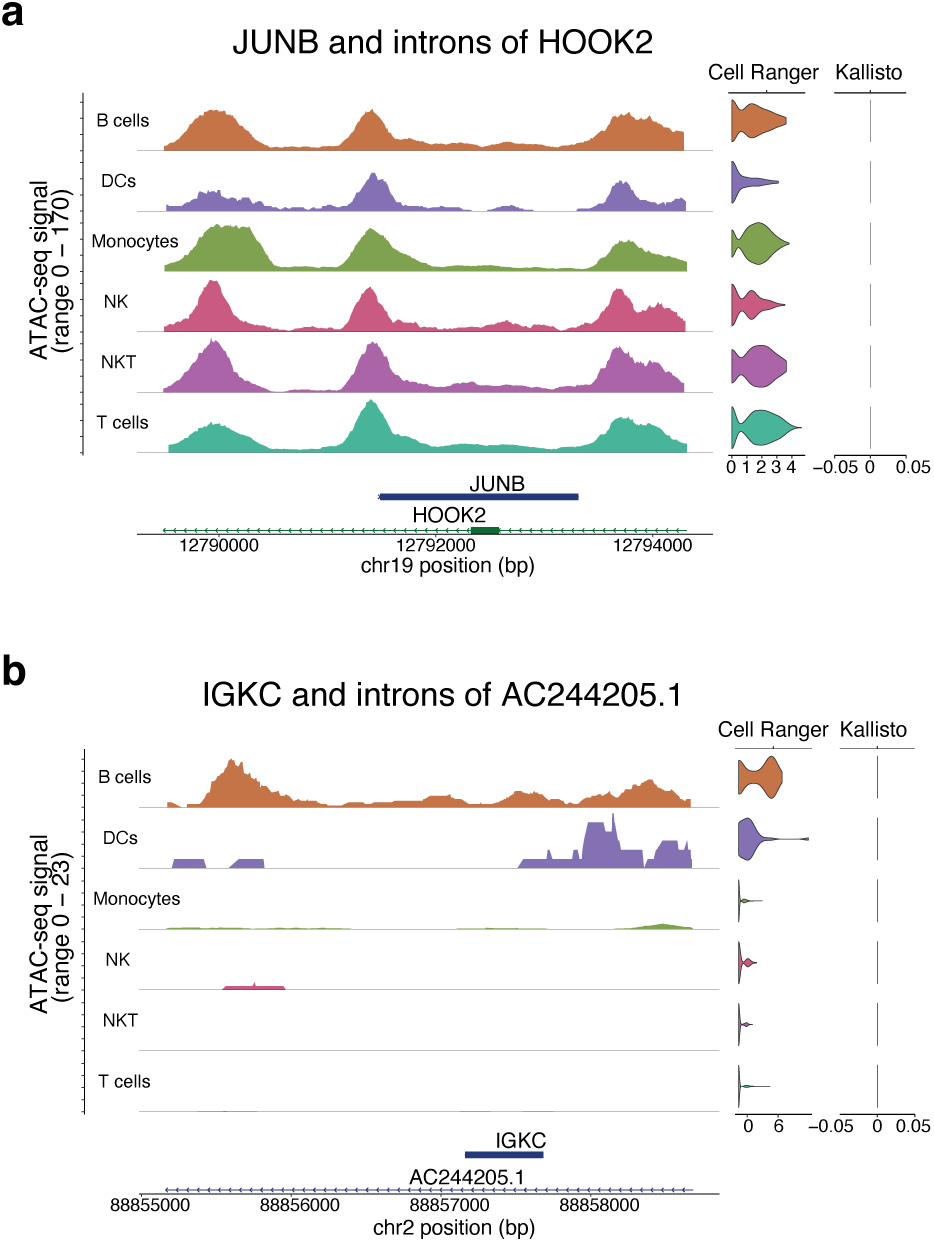
Conflicts with the intronic annotation when an exon of a gene is overlapped with an intronic region of another gene. **a,** ATAC-seq signal and RNA-seq expression, with both Cell Ranger and Kallisto, where part of the mono-exonic *JUNB* is overlapped by an intronic region of *HOOK2*. **b,** ATAC-seq signal and RNA-seq expression, with both Cell Ranger and Kallisto, where the mono-exonic *IGKC* is overlapped by an intronic region of *AC244205.1*.

**Supplementary Data Table 1 (separate file)**

Canonical markers used for distinguishing main cell types.

**Supplementary Data Table 2 (separate file)**

List exclusive lncRNAs that are also hits in public CRISPRi screenings.

**Supplementary Data Table 3 (separate file)**

List of 2,340 lncRNAs that exhibit characteristics of functional lncRNAs in the diverse set of scRNA-seq datasets analyzed.

**Supplementary Data Table 4 (separate file)**

List of 10x Genomics scRNA-seq datasets used.

## Acknowledgements

We thank Dr. Xabier Agirre for providing the KMS-12-BM samples. We particularly acknowledge the patients for their participation and the Biobank of the University of Navarra for its collaboration. This work was supported by grants to: M.He.: H2020 Marie S. Curie IF Action, European Commission, Grant Agreement No. 898356. M.Hu: La Caixa Foundation [LCF/PR/HR21/00176], European Research Council Consolidator 771425 and Worldwide Cancer Research grants 20-0204. P.F: Blanca (0011-1411-2021-000070). E.G. was supported by PhD fellowship from Gobierno de Navarra (0011-0537-2020-000038).

## Author contributions

E.G. designed and performed most analysis. A.M. performed experimental validations. A.A. generated sequencing data from TNBC patients. M.S. collected clinical samples from TNBC patients. P.F. supervised E.G in the scRNA-seq analysis of the TNBC patients. M.Hu. and M.He. conceived the study, supervised the work and obtained funding. E.G., M.Hu. and M.He. wrote the manuscript with input from all authors.

## References

1. Rahman, R. U., Ahmad, I., Sparks, R., Saad, A. Ben & Mullen, A. Singletrome: A method to analyze and enhance the transcriptome with long noncoding RNAs for single cell analysis. doi:10.1101/2022.10.31.514182.

2. Luo, H. et al. Single-cell Long Non-coding RNA Landscape of T Cells in Human Cancer Immunity. Genomics Proteomics Bioinformatics 19, 377– 393 (2021).

3. Zheng, L. L. et al. ColorCells: a database of expression, classification and functions of lncRNAs in single cells. Brief Bioinform 22, 1–11 (2021).

4. Santus, L. et al. Single-cell profiling of lncRNA expression during Ebola virus infection in rhesus macaques. Nature Communications 2023 14:1 14, 1–14 (2023).

5. Statello, L., Guo, C.-J., Chen, L.-L. & Huarte, M. Gene regulation by long non-coding RNAs and its biological functions. Nat Rev Mol Cell Biol doi:10.1038/s41580-020-00315-9.

6. Mattick, J. S. et al. Long non-coding RNAs: definitions, functions, challenges and recommendations. Nature Reviews Molecular Cell Biology 2023 24:6 24, 430–447 (2023).

7. Cabili, M. et al. Integrative annotation of human large intergenic noncoding RNAs reveals global properties and specific subclasses. Genes Dev 25, 1915–1927 (2011).

8. Liu, S. J. et al. Single-cell analysis of long non-coding RNAs in the developing human neocortex. (2016) doi:10.1186/s13059-016-0932-1.

9. Atanasovska, B. et al. A liver-specific long noncoding RNA with a role in cell viability is elevated in human nonalcoholic steatohepatitis. Hepatology 66, 794–808 (2017).

10. Liu, S. J. et al. CRISPRi-based genome-scale identification of functional long noncoding RNA loci in human cells. Science (1979) 355, (2017).

11. Huarte, M. The emerging role of lncRNAs in cancer. Nature Medicine 2015 21:11 21, 1253–1261 (2015).

12. Iyer, M. K. et al. The landscape of long noncoding RNAs in the human transcriptome. Nature Genetics 2015 47:3 47, 199–208 (2015).

13. Frankish, A. et al. GENCODE: reference annotation for the human and mouse genomes in 2023. Nucleic Acids Res 51, D942–D949 (2023).

14. Hezroni, H. et al. Principles of Long Noncoding RNA Evolution Derived from Direct Comparison of Transcriptomes in 17 Species. Cell Rep 11, 1110–1122 (2015).

15. Uszczynska-Ratajczak, B., Lagarde, J., Frankish, A., Guigó, R. & Johnson, R. Towards a complete map of the human long non-coding RNA transcriptome. Nature Reviews Genetics 2018 19:9 19, 535–548 (2018).

16. Aldridge, S. & Teichmann, S. A. Single cell transcriptomics comes of age. Nature Communications 2020 11:1 11, 1–4 (2020).

17. Zeisel, A. et al. Molecular Architecture of the Mouse Nervous System. Cell 174, 999–1014.e22 (2018).

18. Prescott, S. L., Umans, B. D., Williams, E. K., Brust, R. D. & Liberles, S. D. An Airway Protection Program Revealed by Sweeping Genetic Control of Vagal Afferents. Cell 181, 574–589.e14 (2020).

19. La Manno, G. et al. RNA velocity of single cells. Nature 2018 560:7719 560, 494–498 (2018).

20. Macosko, E. Z. et al. Highly Parallel Genome-wide Expression Profiling of Individual Cells Using Nanoliter Droplets. Cell 161, 1202–1214 (2015).

21. Klein, A. M. et al. Droplet Barcoding for Single-Cell Transcriptomics Applied to Embryonic Stem Cells. Cell 161, 1187–1201 (2015).

22. Zheng, G. X. Y. et al. Massively parallel digital transcriptional profiling of single cells. Nature Communications 2017 8:1 8, 1–12 (2017).

23. Svensson, V., Vento-Tormo, R. & Teichmann, S. A. Exponential scaling of single-cell RNA-seq in the past decade. Nature Protocols 2018 13:4 13, 599–604 (2018).

24. Papalexi, E. & Satija, R. Single-cell RNA sequencing to explore immune cell heterogeneity. Nature Reviews Immunology 2017 18:1 18, 35–45 (2017).

25. Hwang, B., Lee, J. H. & Bang, D. Single-cell RNA sequencing technologies and bioinformatics pipelines. Exp Mol Med 53, 1005–1005 (2021).

26. Luecken, M. D. & Theis, F. J. Current best practices in single-cell RNA-seq analysis: a tutorial. Mol Syst Biol 15, e8746 (2019).

27. You, Y. et al. Benchmarking UMI-based single-cell RNA-seq preprocessing workflows. Genome Biol 22, 339 (2021).

28. Dobin, A. et al. STAR: ultrafast universal RNA-seq aligner. Bioinformatics 29, 15–21 (2013).

29. Kaminow, B., Yunusov, D. & Dobin, A. STARsolo: accurate, fast and versatile mapping/quantification of single-cell and single-nucleus RNA-seq data. bioRxiv 2021.05.05.442755 (2021) doi:10.1101/2021.05.05.442755.

30. Bray, N. L., Pimentel, H., Melsted, P. & Pachter, L. Near-optimal probabilistic RNA-seq quantification. Nature Biotechnology 2016 34:5 34, 525–527 (2016).

31. Patro, R., Duggal, G., Love, M. I., Irizarry, R. A. & Kingsford, C. Salmon provides fast and bias-aware quantification of transcript expression. Nature Methods 2017 14:4 14, 417–419 (2017).

32. Melsted, P. et al. Modular, efficient and constant-memory single-cell RNA-seq preprocessing. Nature Biotechnology 2021 39:7 39, 813–818 (2021).

33. Melsted, P., Ntranos, V. & Pachter, L. The barcode, UMI, set format and BUStools. Bioinformatics 35, 4472–4473 (2019).

34. Srivastava, A., Malik, L., Smith, T., Sudbery, I. & Patro, R. Alevin efficiently estimates accurate gene abundances from dscRNA-seq data. Genome Biol 20, 1–16 (2019).

35. Vieth, B., Parekh, S., Ziegenhain, C., Enard, W. & Hellmann, I. A systematic evaluation of single cell RNA-seq analysis pipelines. Nature Communications 2019 10:1 10, 1–11 (2019).

36. Brüning, R. S., Tombor, L., Schulz, M. H., Dimmeler, S. & John, D. Comparative analysis of common alignment tools for single-cell RNA sequencing. Gigascience 11, (2022).

37. Du, Y., Huang, Q., Arisdakessian, C. & Garmire, L. X. Evaluation of STAR and Kallisto on Single Cell RNA-Seq Data Alignment. G3 Genes|Genomes|Genetics 10, 1775–1783 (2020).

38. Derrien, T. et al. The GENCODE v7 catalog of human long noncoding RNAs: Analysis of their gene structure, evolution, and expression. doi:10.1101/gr.132159.111.

39. Zheng, H., Brennan, K., Hernaez, M. & Gevaert, O. Benchmark of long non-coding RNA quantification for RNA sequencing of cancer samples. 8, 1–13 (2019).

40. 1k Brain Cells from an E18 Mouse (v3 chemistry) - 10x Genomics. https://www.10xgenomics.com/resources/datasets/1-k-brain-cells-from-an-e-18-mouse-v-3-chemistry-3-standard-3-0-0.

41. PBMCs from a Healthy Donor: Whole Transcriptome Analysis - 10x Genomics. https://www.10xgenomics.com/resources/datasets/pbm-cs-from-a-healthy-donor-whole-transcriptome-analysis-3-1-standard-4-0-0.

42. He, D. et al. Alevin-fry unlocks rapid, accurate and memory-frugal quantification of single-cell RNA-seq data. Nature Methods 2022 19:3 19, 316–322 (2022).

43. Fawkner-Corbett, D. et al. Spatiotemporal analysis of human intestinal development at single-cell resolution ll Spatiotemporal analysis of human intestinal development at single-cell resolution. Cell 184, 810–826 (2021).

44. Schupp, J. C. et al. Integrated Single-Cell Atlas of Endothelial Cells of the Human Lung. Circulation 144, 286–302 (2021).

45. Habermann, A. C. et al. Single-cell RNA sequencing reveals profibrotic roles of distinct epithelial and mesenchymal lineages in pulmonary fibrosis. Sci Adv 6, (2020).

46. 10k Mouse PBMCs Multiplexed, 2 CMOs - 10x Genomics. https://www.10xgenomics.com/resources/datasets/10-k-mouse-pbm-cs-multiplexed-2-cm-os-3-1-standard-6-0-0.

47. 5k Peripheral Blood Mononuclear Cells (PBMCs) from a Healthy Donor (Next GEM) - 10x Genomics. https://www.10xgenomics.com/resources/datasets/5-k-peripheral-blood-mononuclear-cells-pbm-cs-from-a-healthy-donor-next-gem-3-1-standard-3-0-2.

48. PBMC from a Healthy Donor - Granulocytes Removed Through Cell Sorting (3k) - 10x Genomics. https://www.10xgenomics.com/resources/datasets/pbmc-from-a-healthy-donor-granulocytes-removed-through-cell-sorting-3-k-1-standard-2-0-0.

49. Kirk, J. M. et al. Functional classification of long non-coding RNAs by k-mer content. Nature Genetics 2018 50:10 50, 1474–1482 (2018).

50. Wu, S. Z. et al. A single-cell and spatially resolved atlas of human breast cancers. Nature Genetics 2021 53:9 53, 1334–1347 (2021).

51. Namba, M. et al. Establishment of five human myeloma cell lines. In Vitro Cellular & Developmental Biology 25, 723–729 (1989).

52. Edwards, J. C. W. & Cambridge, G. B-cell targeting in rheumatoid arthritis and other autoimmune diseases. Nature Reviews Immunology 2006 6:5 6, 394–403 (2006).

53. Jourdan, M. et al. An in vitro model of differentiation of memory B cells into plasmablasts and plasma cells including detailed phenotypic and molecular characterization. Blood 114, 5173–5181 (2009).

54. Bitar, M. et al. Redefining normal breast cell populations using long noncoding RNAs. Nucleic Acids Res (2023) doi:10.1093/nar/gkad339.

55. Srivastava, A. et al. Alignment and mapping methodology influence transcript abundance estimation. Genome Biol 21, 1–29 (2020).

56. Shainer, I. & Stemmer, M. Choice of pre-processing pipeline influences clustering quality of scRNA-seq datasets. BMC Genomics 22, (2021).

57. Wang, H. et al. Selective effects of protein 4.1N deficiency on neuroendocrine and reproductive systems. Scientific Reports 2020 10:1 10, 1–14 (2020).

58. Kim, A. C., Van Huffel, C., Lutchman, M. & Chishti, A. H. Radiation Hybrid Mapping ofEPB41L1,a Novel Protein 4.1 Homologue, to Human Chromosome 20q11.2–q12. Genomics 49, 165–166 (1998).

59. Hjörleifsson, K. E., Sullivan, D. K., Holley, G., Melsted, P. & Pachter, L. Accurate quantification of single-nucleus and single-cell RNA-seq transcripts. doi:10.1101/2022.12.02.518832.

60. He, D., Soneson, C. & Patro, R. Understanding and evaluating ambiguity in single-cell and single-nucleus RNA-sequencing. bioRxiv 2023.01.04.522742 (2023) doi:10.1101/2023.01.04.522742.

61. Pool, A. H., Poldsam, H., Chen, S., Thomson, M. & Oka, Y. Recovery of missing single-cell RNA-sequencing data with optimized transcriptomic references. Nature Methods 2023 20:10 20, 1506–1515 (2023).

62. Chakraborty, S. et al. Harnessing the tissue and plasma lncRNA-peptidome to discover peptide-based cancer biomarkers. Scientific Reports 2019 9:1 9, 1–17 (2019).

63. Goyal, B. et al. Diagnostic, prognostic, and therapeutic significance of long non-coding RNA MALAT1 in cancer. BBA-Reviews on Cancer 1875, 188502 (2021).

64. SC5P-R2 sequencing · Issue #226 · pachterlab/kallisto. https://github.com/pachterlab/kallisto/issues/226.

65. Selective Alignment. https://combine-lab.github.io/alevin-tutorial/2019/selective-alignment/.

66. Amezquita, R. A. et al. Orchestrating single-cell analysis with Bioconductor. Nature Methods 2019 17:2 17, 137–145 (2019).

67. Lun, A. T. L. et al. EmptyDrops: Distinguishing cells from empty droplets in droplet-based single-cell RNA sequencing data. Genome Biol 20, 1–9 (2019).

68. Germain, P. L., Lun, A., Macnair, W. & Robinson, M. D. Doublet identification in single-cell sequencing data using scDblFinder. F1000Research 2021 10:979 10, 979 (2021).

69. LTLA/scuttle: Clone of the Bioconductor repository for the scuttle package. https://github.com/LTLA/scuttle/.

70. McCarthy, D. J., Campbell, K. R., Lun, A. T. L. & Wills, Q. F. Scater: pre-processing, quality control, normalization and visualization of single-cell RNA-seq data in R. Bioinformatics 33, 1179–1186 (2017).

71. Lun, A. T. et al. A step-by-step workflow for low-level analysis of single-cell RNA-seq data with Bioconductor. F1000Research 2016 5:2122 5, 2122 (2016).

72. Network Analysis and Visualization [R package igraph version 1.5.1]. (2023).

73. igraph – Network analysis software. https://igraph.org/.

74. Goyal, M. et al. JIND: joint integration and discrimination for automated single-cell annotation. Bioinformatics 38, 2488–2495 (2022).

75. Joint RNA and ATAC analysis: 10x multiomic • Signac. https://stuartlab.org/signac/articles/pbmc_multiomic.

76. Stuart, T., Srivastava, A., Madad, S., Lareau, C. A. & Satija, R. Single-cell chromatin state analysis with Signac. Nature Methods 2021 18:11 18, 1333–1341 (2021).

77. RepeatMasker Home Page. https://www.repeatmasker.org/.

78. CalabreseLab/seekr: A library for counting small kmer frequencies in nucleotide sequences. https://github.com/CalabreseLab/seekr.

79. Camargo, A. P., Vasconcelos, A. A., Fiamenghi, M. B., Pereira, G. A. G. & Carazzolle, M. F. tspex: a tissue-specificity calculator for gene expression data. 1–7 (2020) doi:10.21203/RS.3.RS-51998/V1.

80. Hao, Y. et al. Integrated analysis of multimodal single-cell data. Cell 184, 3573–3587.e29 (2021).

81. Conway, J. R., Lex, A. & Gehlenborg, N. UpSetR: an R package for the visualization of intersecting sets and their properties. Bioinformatics 33, 2938–2940 (2017).

